# Neuropilin-2 functions as a co-inhibitory receptor to regulate antigen-induced inflammation and allograft rejection

**DOI:** 10.1101/2025.04.17.648233

**Authors:** Johannes Wedel, Nora Kochupurakkal, Sek Won Kong, Sayantan Bose, Ji-Won Lee, Madeline Maslyar, Bayan Alsairafi, Kayla Macleod, Kaifeng Liu, Hengcheng Zhang, Masaki Komatsu, Hironao Nakayama, Diane R. Bielenberg, David M. Briscoe

**Author notes:** **Address for Correspondence:** David M. Briscoe, M.D., Transplant Research Program, Division of Nephrology, Boston Children’s Hospital, 300 Longwood Ave, Boston, MA 02115. **Conflict-of-interest statement:** All authors have no conflicts of interest.

## Abstract

Co-inhibitory receptors function as central modulators of the immune response to resolve T effector activation and/or to sustain immune homeostasis. Here, using humanized SCID mice, we found that NRP2 is inducible on late effector and exhausted subsets of human CD4^+^ T cells and that it is co-expressed with established co-inhibitory molecules including PD-1, CTLA4, TIGIT, LAG3 and TIM3. In murine models, we also found that NRP2 is expressed on effector memory CD4^+^ T cells with an exhausted phenotype and that it functions as a key co-inhibitory molecule. Knockout of NRP2 resulted in hyperactive CD4^+^ T cell responses, and enhanced inflammation in delayed type hypersensitivity and transplantation models. Following cardiac transplantation, allograft rejection and graft failure was accelerated in global as well as CD4^+^ T cell-specific knockout recipients, and enhanced alloimmunity was dependent on NRP2 expression on CD4^+^ T effectors, and not on CD4^+^Foxp3^+^ T regulatory cells. Also, knockout T regulatory cells were found to be as efficient as wild type cells in the suppression of effector responses in vitro and in vivo. These collective findings identify NRP2 as a novel coinhibitory receptor and demonstrate that its expression on CD4^+^ T effector cells is of great functional importance in immunity.

## Introduction

T cell activation is intrinsically regulated and self-limited to prevent chronic inflammation and autoimmunity. One such regulatory mechanism is called T cell exhaustion (Texh cells, also called T cell dysfunction), characterized by reduced T effector function including loss of cytokine production and reduced proliferation rates (1). Differentiation into Texh is reported to be in-part dependent on the induced expression of co-inhibitory receptors including PD-1, CTLA4, TIGIT, LAG3, TIM-3 and BTLA (1–4). These co-inhibitory receptors collectively function to suppress ongoing cell-mediated immune activation which ultimately serves to resolve the inflammatory response. Augmentation of Texh has multiple regulatory effects on the immune response including an established effect to limit alloimmunity and prolong graft survival following transplantation (5). In contrast, blockade of individual co-inhibitory receptors may reverse Texh for example to enhance tumor immune responses (6–12). Thus, understanding the cellular basis for Texh/T cell dysfunction has broad clinical implications (5, 13–15).

The neuropilin (NRP) receptors NRP1 and NRP2 are type I transmembrane glycoproteins that were initially identified as chemorepulsive axonal guidance receptors (16–19). These receptors bind multiple ligands including class III semaphorins (SEMA3), vascular endothelial growth factor (VEGF) -A and -C, and transforming growth factor beta (TGF-β) (17–24). They are expressed by multiple cell types and function in a broad range of biological processes including cytoskeletal stability, migration, angiogenesis, and cell growth (25–29). NRP1 also functions to regulate cell-mediated immune responses via its expression on both T cells and antigen presenting cells (APC) (30, 31). Indeed, a dominant biological effect of NRP1 relates to its expression on CD4^+^ T regulatory cells (Tregs) where it augments immunoregulation and lineage stability (32–36). Deletion of NRP1 on CD4^+^ T cells is reported to increase disease severity in models of experimental autoimmune encephalitis (37) and colitis (34). Also, NRP1 is reported to regulate CD8^+^ T cell memory responses in association with anti-tumor immunity (38, 39).

In this study, we identify NRP2 on subsets of human and mouse immune cells and we show that it is inducible on antigen-activated CD4^+^ T cells, most notably on late effector and Texh subsets. Using NRP2 knock-out mice, we find that it functions to regulate CD4^+^ T cell activation in vitro as well as cell-mediated immune responses following vaccination and transplantation in vivo. We also find that knockout of NRP2 on CD4^+^ Teffs, but not on CD4^+^Foxp3^+^ Tregs is associated with accelerated rejection and enhanced alloimmunity following cardiac transplantation. Furthermore, in a skin transplantation model, we find that NRP2 is redundant on Treg cells, as deletion does not alter their immune suppressive function in vitro. Overall, our findings demonstrate that NRP2 is a co-inhibitory receptor on CD4^+^ T cells and that it functions to regulate antigen-activated and effector immune responses most notably following transplantation.

## Results

### NRP2 is expressed by multiple immune cell subsets and is inducible on subpopulations of CD4^+^ T cells in the course of an immune response

We initially profiled NRP2 expression on human peripheral blood mononuclear cells (PBMC). As illustrated in Figure 1A-B, it is expressed on several cell types including CD4^+^ and CD8^+^ T cells, CD19^+^ B cells and CD11c^+^ antigen presenting cells (APCs). Moreover, we consistently we found a most notable expression pattern on a subpopulation of CD4^+^ T cells by flow cytometry (Figure 1A, left panel and Supplemental Figure 1). Expression was prominent on isolated CD4^+^ T cells by immunofluorescence staining and confocal microscopy (Figure 1C). Also, we find that the level of NRP2 expression on distinct subpopulation(s) of CD4^+^ T cells is sustained following mitogen activation (Figure 1D-E).

**Figure 1:**
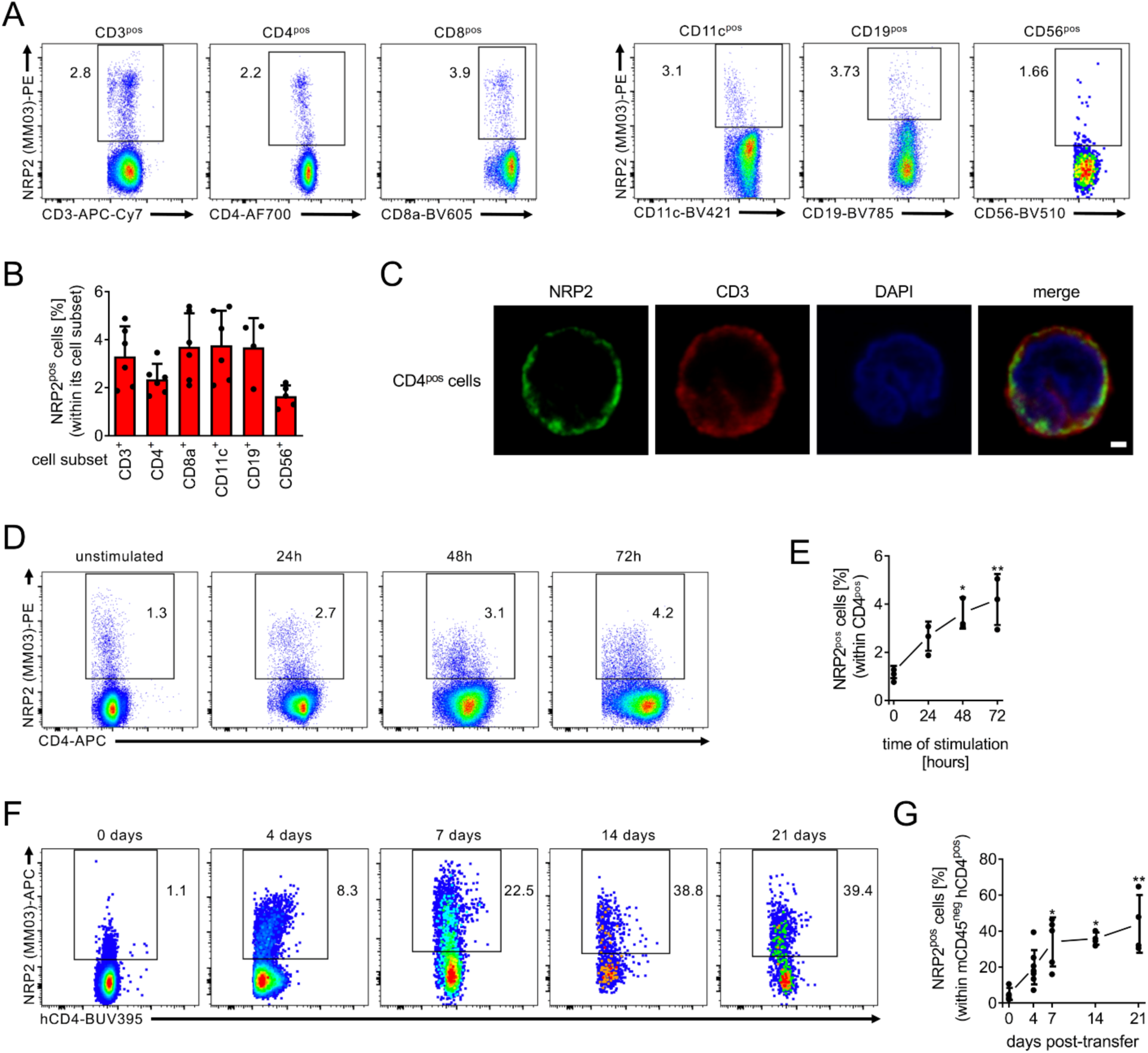
Inducible NRP2 expression on distinct subset(s) of human CD4^+^ T cells. **(A)** Representative dot plots, and **(B)** a summary of six independent flow cytometric analyses (mean ± SD) of NRP2 staining of freshly isolated human PBMC. **(C)** Representative cytospin of negatively isolated human CD4^+^ T cells stained for NRP2 (clone MM03; green) and CD3 (red), and imaged by confocal microscopy. Representative images of four independent experiments showing a NRP2-expressing CD4^+^ cell at high power magnification (bar represent 1 µm). **(D)** Representative dot plots of NRP2 expression on human CD4^+^ T cells cultured in the presence of phytohemagglutinin (PHA; 3 µg/ml for up to 72 hours) and evaluated by flow cytometry. **(E)** Line graph summarizing three independent experiments (mean ± SD; Friedman’s test with Dunn’s multiple comparison, * *P*<0.05, ** *P*<0.01). **(F)** NRP2 expression by flow cytometry on human CD4^+^ T cells within splenocytes of huSCID mice at selected time intervals following humanization. Dot plots are gated on human CD4^+^ cells (murine CD45^neg^). **(G)** Line graph illustrating changes in the expression of NRP2 on CD4^+^ T cells over a 21-day period following humanization of huSCID mice (n=4-8 independent experiments per time-point; mean ± SD; Kruskal-Wallis test with Dunn’s multiple comparison, * *P*<0.05, ** *P*<0.01).

We next transferred PBMC into lymphopenic SCID-beige mice, harvested the spleen and evaluated expression on transferred cells as a time course for up to 21 days (Figure 1F-G and Supplemental Figure 2). By flow cytometry, we find a marked induction in NRP2 expression on CD4^+^ T cells (Figure 1F-G) as well as on CD8^+^ T cells, CD11c^+^ APCs and CD19^+^ B cells (Supplemental Figure 2D-G). Expression within the CD4^+^ population increased from <3% cells at baseline to over 30% cells on day 14 and up to ∼45% cells over the 21-day time course (Figure 1G). Collectively, these findings indicate that NRP2 is expressed on multiple immune cell lineages but is markedly inducible on distinct subset(s) of CD4^+^ T cells following activation in vivo.

### Co-expression of NRP2 with multiple co-inhibitory receptors on CD4^+^ T cells

To identify the subpopulation(s) of CD4^+^ cells that express NRP2, we performed cellular indexing of transcriptomes and epitopes (CITE) sequencing (scRNAseq) on CD4^+^ T cells that were isolated on day 7 post transfer in the huSCID model (Figure 1F and Supplemental Figure 3A). t-SNE dimensional reduction and clustering was performed (Figure 2A), and NRP2 positivity was evaluated using two NRP2 antibodies (Supplemental Figure 3B). We find that NRP2 concentrated in two clusters belonging to PD-1^+^TIM-3^+^EOMES^neg^ late-stage T effectors and PD-1^+^TIM- 3^+^EOMES^pos^ exhausted subsets (Figure 2B and Supplemental Figure 3C). Pseudotime trajectory analysis (Figure 2C-D) demonstrated that NRP2 expression starts within resting CD4^+^ T cells and progresses through activated and late-stage Teffs and ultimately within Texh cell subsets (Figure 2C). Heatmap analysis along the pseudotime further indicate that NRP2 expression is initiated during Teff cell activation and peaks during later stages of Teff differentiation in association with exhaustion (Figure 2D). Finally, we performed flow cytometric analysis at multiple time points following transfer of PBMC in the huSCID mouse and find that NRP2 co-localizes with established co-inhibitory molecules including PD-1, CTLA4, TIGIT, LAG-3 and TIM-3 further confirming that it is expressed on phenotypic Texh subsets (Figure 2E and Supplemental Figure 3D-E). Consistent with this phenotype, NRP2^pos^ subsets had a significantly lower proliferation rate (as assessed by BrdU incorporation) and produced minimal IFN-γ compared to NRP2^neg^ CD4^+^ T cells (Figure 2F-G). These findings are most suggestive that NRP2^pos^ cells are both phenotypically and functionally exhausted. Finally, expression of CD69, CD38 and HLA-DR were not associated with NRP2 positivity indicating that it is not a general marker of activation (Figure 2H-J). Collectively, these data identify NRP2 on late-stage effector and Texh CD4^+^ T cell subsets.

**Figure 2:**
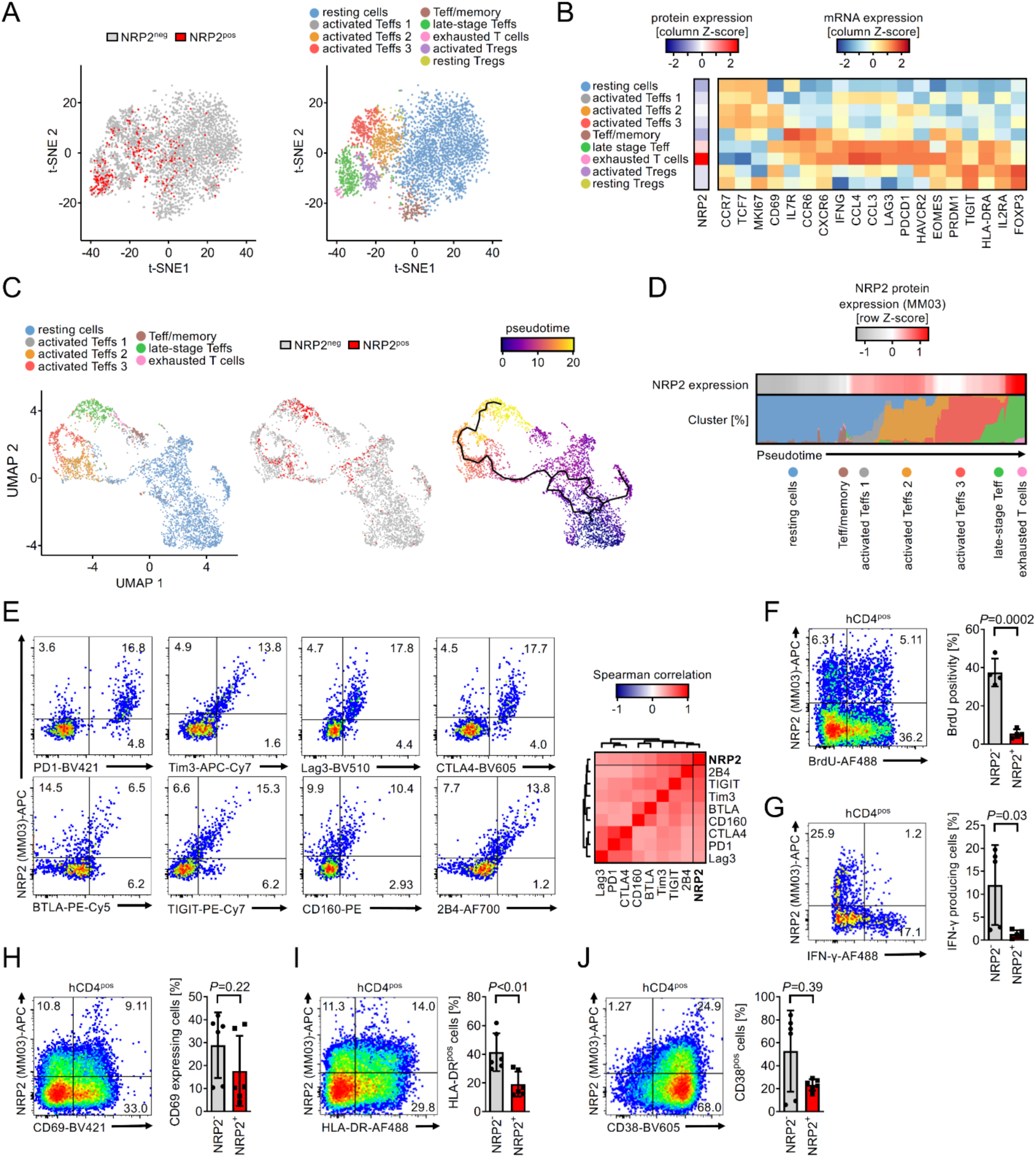
Patterns of expression of NRP2 on late-stage effector and exhausted CD4^+^ T cells. Human CD4^+^ T cells were isolated from the spleens of huSCID mice on day 7 following humanization; the cells were stained with DNA- and fluorochrome-conjugated anti-NRP2 antibodies, and subsequently sorted by FACS for proteogenomic analysis using CITE-seq. **(A)** t-SNE plots depicting NRP2 protein expression (left) and CD4^+^ T cell subset clusters based on transcriptome (right). **(B)** Heatmap of cluster-defining transcripts for Panel A. **(C)** UMAP plots with embedding of Teff populations (excluding Treg clusters) with color coding of each cluster (left), color coding of NRP2 protein expression (middle) and color coding of the calculated pseudotime (right; solid black line represents the pseudotime trajectory). **(D)** Color coding to represent the level of NRP2 protein expression over the pseudotime in Panel C. **(E)** Representative flow cytometric expression of coinhibitory molecules on isolated CD4^+^ T cells (left) and a heatmap illustrating the mean spearman’s rank correlation coefficient between NRP2 and each coinhibitory receptor in four independent experiments (right). **(F)** huSCID mice were pulsed with BrdU on day 4-7 post-transfer and spleens were harvested on day 7; the proliferation of human CD4^+^ T cells was assessed by intracellular BrdU staining using flow cytometry. A representative dot plot (left) and a bar graph summary of BrdU positivity in n=4 mice (right) are depicted (mean ± SD; unpaired t-test). **(G)** Intracellular IFN-γ staining of human CD4^+^ cells (day 7 post transfer). A representative dot plot (left) and a summary of IFN-γ producing cells in n=5 mice (right) are depicted (mean ± SD; Mann Whitney test). **(H-J)** Expression of the activation markers CD69, HLA-DR and CD38 on human CD4^+^ cells (day 7). Representative dot plots (left) and a summary of expression in n=6 mice (right) are depicted (mean ± SD; H-I: unpaired t-test, J: Mann Whitney test).

### NRP2-expressing CD4^+^ T effector cells are enriched with genes associated with T cell dysfunction

Bulk RNA-sequencing of FACS-sorted NRP2^pos^ and NRP2^neg^ CD4^+^ T cells revealed that NRP2^pos^ subsets have a unique transcriptomic signature that differentiates them from NRP2^neg^ cells (Figure 3A and Supplemental Figure 4). Furthermore, this signature is sustained following activation with PHA (Figure 3A). In unstimulated conditions, differences among NRP2^pos^ and NRP2^neg^ cells include subclusters of transcripts for costimulatory molecules, chemokine receptors and kynurenine system molecules (Figure 3B-C). In the presence of mitogen, differences also include transcripts for select cytokines including IL-6 (Figure 3C). Additionally, gene set enrichment analyses indicated that unstimulated NRP2^pos^ CD4^+^ T cell subsets express transcripts associated with exhaustion/dysfunction (Figure 3D), and this transcriptome is sustained following mitogen activation (data not shown). Finally, computed intrinsic differences in signaling pathway activities were minimal in unstimulated NRP2^pos^ and NRP2^neg^ CD4^+^ subsets (Supplemental Figure 4C) whereas mitogen activation resulted in identifiable effects of NRP2 on pathways that include NF- κB, STAT3 and cell cycle associated signaling (Supplemental Figure 4C). Collectively, these findings indicate that NRP2^pos^ CD4^+^ T cells possess a unique transcriptome, and thus are likely a distinct subset.

**Figure 3:**
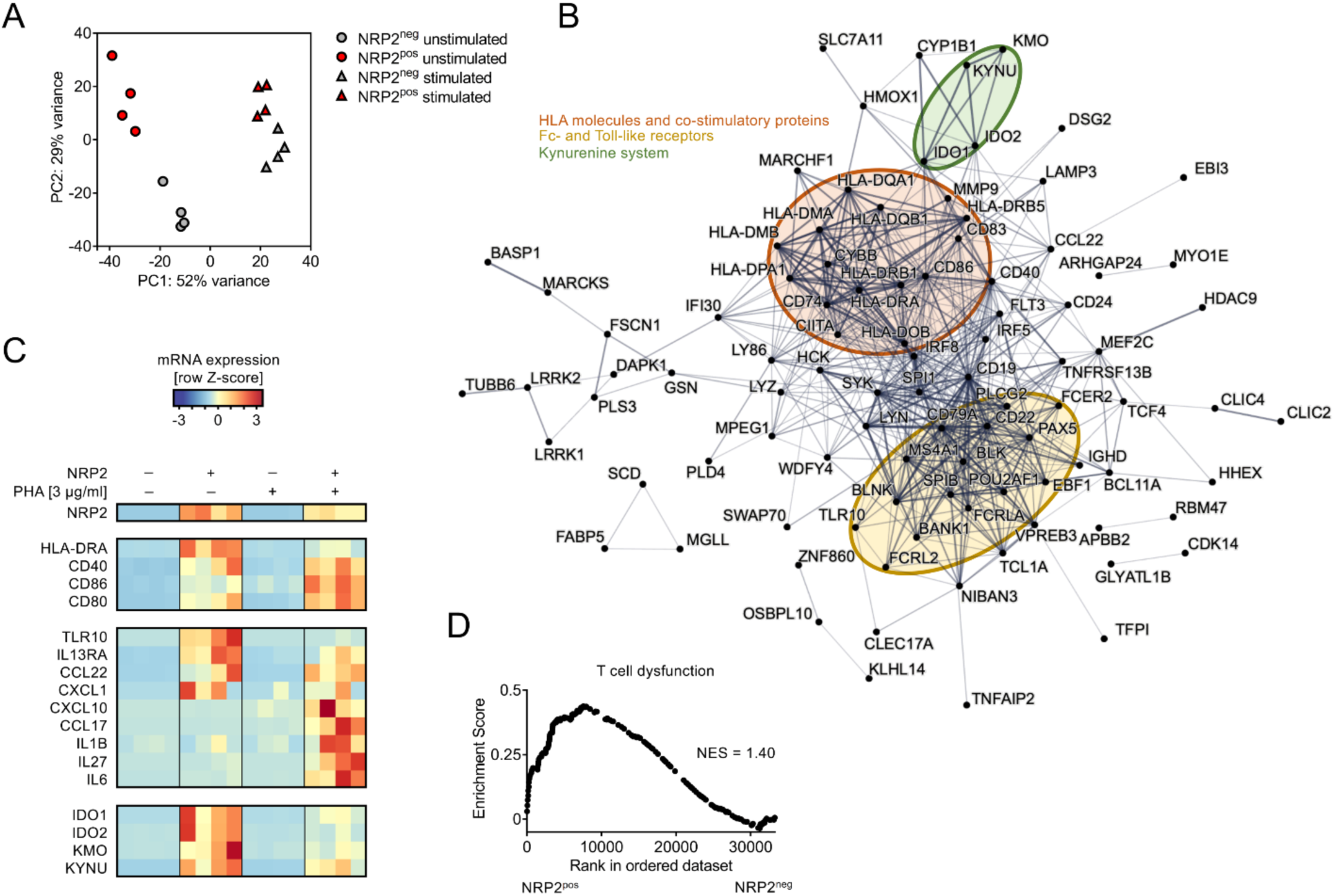
NRP2^pos^ CD4^+^ T cells have a distinct transcriptional profile. Pooled populations of human CD4^+^ T cells were stimulated with PHA (3 µg/ml) for 16 hours in vitro and NRP2^pos^ and NRP2^neg^ cells were sorted by flow cytometry and subjected to transcriptomic analysis. **(A)** Principal component analysis of unstimulated and stimulated subsets. **(B)** Protein-Protein interaction analysis using NRP2 co-regulated transcripts (≥2.5 log fold change and *Padj* < 10^-10^ between unstimulated NRP2^pos^ and NRP2^neg^ cells shown in Panel A). Subnetwork nodes are highlighted in color. **(C)** Heatmap-depicting transcripts identified in protein interaction network analyses in Panel B and additional transcripts that are upregulated in NRP2^pos^ cells following PHA stimulation. **(D)** Enrichment for genes associated with dysfunctional T cells (90) using Gene Set Enrichment Analysis. Ranked in order for NRP2^pos^ to NRP2^neg^ cells in unstimulated conditions (NES, Normalized Enrichment Score).

### NRP2 serves as a co-inhibitory receptor and regulates CD4^+^ T effector function

Similar to human cells (Figure 1), we find that NRP2 is expressed on a small but distinct subset of murine CD4^+^ T effector cells, and that it is inducible following mitogen-activation in vitro and allopriming in vivo (Figure 4A-E and Supplemental Figures 5-6). We also adoptively transferred ovalbumin-specific T cell receptor transgenic CD4^+^ T cells (OT-II, CD45.2) into CD45.1 hosts and profiled NRP2 expression on CD45.1 and CD45.2 cells following immunization with ovalbumin. As illustrated in Figure 4G-H, we find a 3-fold induction in NRP2 mRNA expression (using PrimeFlow cytometry) on antigen-specific OT-II vs. host CD4^+^ T cells and, consistent with our findings in Figure 2, murine NRP2^hi^ CD4^+^ T cells co-express PD-1 and TIM-3 (Figure 4I-J). Also, as expected, antigen-specific NRP-2^hi^ CD4^+^ T cells were CD44^high^CD62L^low^ consistent with an effector/memory phenotype (data not shown).

**Figure 4:**
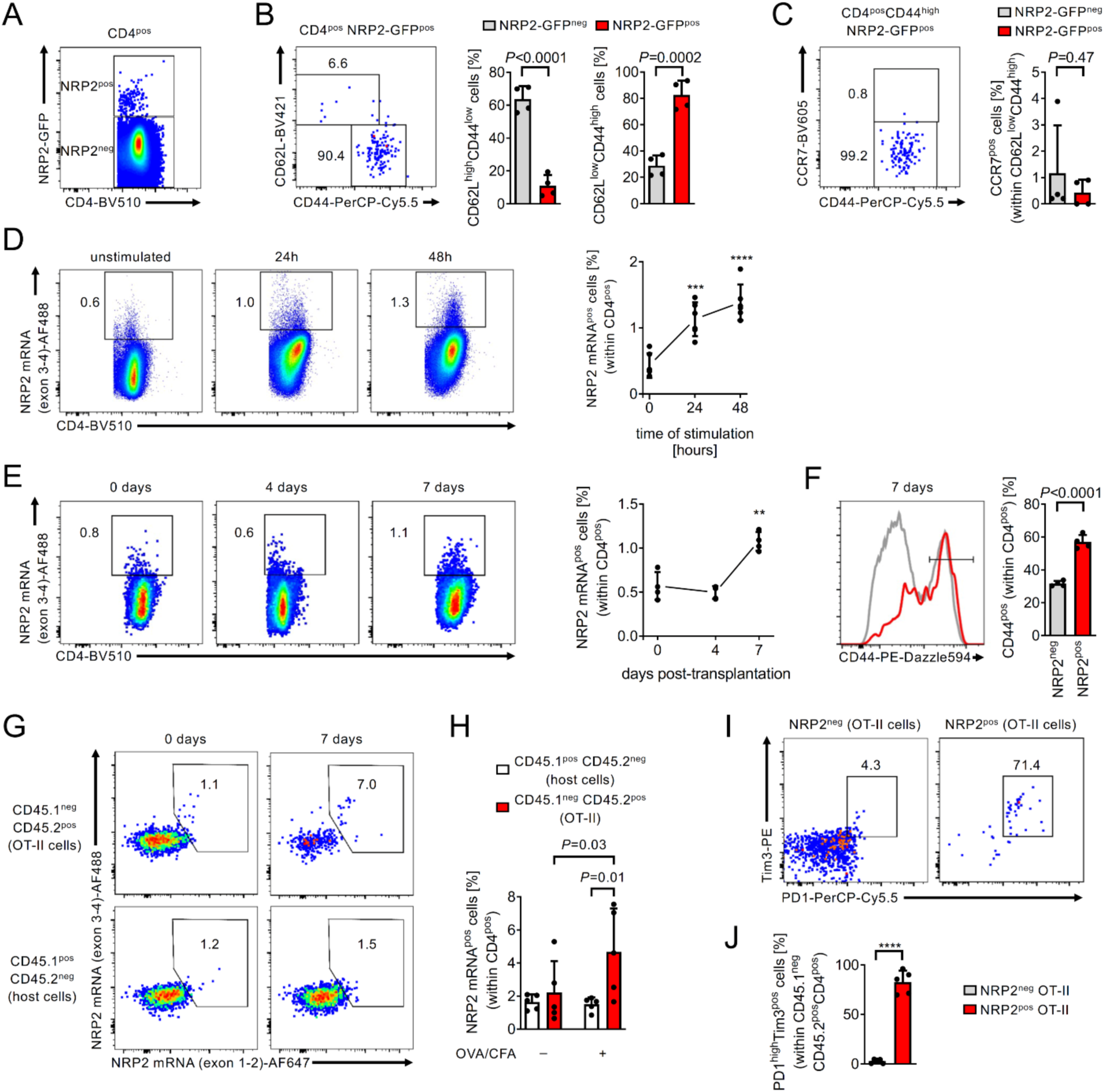
Patterns of NRP2 expression on murine CD4^+^ T cells following activation. **(A)** Representative dot plot of NRP2-GFP expression within CD4^+^ splenocytes of NRP2^lox/lox^ mice (n=4 independent experiments; see Supplemental Figure 5A-D). **(B-C)** Representative dot plots (left) and a summary of n=4 independent experiments comparing the phenotype of NRP2-GFP^pos^ to NRP2-GFP^neg^ CD4^+^ splenocytes. **(D)** NRP2 mRNA expression by flow cytometry on isolated murine CD4^+^ T cells stimulated with 1 µg/ml anti-CD3 for up to 48h in vitro. Dot plots are gated on CD4^+^ cells. Line graph illustrates changes in the expression of NRP2 mRNA in CD4^+^ T cells over 48 hours of in vitro stimulation (n=6 independent experiments; mean ± SD; Kruskal-Wallis test with Dunn’s multiple comparison, *** *P*<0.001; **** *P*<0.0001 vs. unstimulated). **(E-F)** Fully MHC mismatched Balb/c donor hearts were transplanted into C57BL/6 recipients. Splenocytes were isolated 4-7 days post-transplant and the frequency NRP2 mRNA expressing CD4^+^ T cells were evaluated by flow cytometry. Non-transplanted C57BL/6 mice were included to illustrate NRP2 mRNA expression pre-transplant (day 0). **(E)** Dot plots are gated on CD4^+^ cells. Line graph illustrates changes in the expression of NRP2 mRNA in CD4^+^ T cells pre- and up to 7 days post-transplant (n=4 mice per time-point; mean ± SD; Kruskal-Wallis test with Dunn’s multiple comparison, \*\**P*<0.01 vs. day 0). **(F)** CD44 expression of NRP2^neg^ and NRP2^pos^ CD4^+^ T cells isolated 7 days post-transplant. Line graph summarizes n=4 experiments (paired t-test). **(G- J)** 2.5 x 10^6^ CD4^+^ T cells from CD45.2^pos^ OT-II mice were adoptively transferred into congenic CD45.1^+^ hosts by tail vein injection. Host mice were immunized subcutaneously with ovalbumin (50 µg) in complete Freud’s adjuvant (CFA), and the phenotype of antigen-specific OT-II and host CD4^+^ T cells were assessed 7 days post-immunization by flow cytometry. **(G)** Representative dot plots of CD45.2^pos^ OT-II (top) and CD45.1^pos^ host (bottom) CD4^+^ T cells are shown. **(H)** Bar graphs represent mean NRP2 positivity within CD4^+^ T cells ± SD of n=5/condition (Two-way ANOVA with Fisher’s Least Significant Difference test). **(I)** Representative dot plots illustrate PD1 and Tim3 expression of NRP2^pos^ (right) and NRP2^neg^ (left) OT-II CD4^+^ T cells. **(H)** Bar graphs summarize the mean ± SD of PD1^+^Tim3^+^ cells of n=5/condition (paired t-test).

To next assess the function of NRP2, we generated global NRP2 knockout (KO, NRP2^-/-^), conditional CD4-specific NRP2 KO (ΔNRP2-CD4) and Foxp3-specific NRP2 KO (ΔNRP2- Foxp3) mice. All transgenic mice were viable for >6 months, remained healthy and did not develop clinical signs of autoimmunity. Furthermore, phenotyping of T cells from thymus, lymph node and spleen demonstrated normal T cell development and subset survival (Supplemental Figure 7). However, CD4^+^ T cells isolated from global NRP2^-/-^ KO and ΔNRP2-CD4 KO mice were hyperproliferative and produced increased IL-2, IFN-γ and IL-17 following mitogenic activation in vitro (Figure 5A-C). Furthermore, culture of CD4^+^ T cells from NRP2^-/-^ or ΔNRP2-CD4 mice in T cell differentiation media resulted in enhanced Th1 and Th2 responses and a trend towards increased Th17 responses vs. WT cells (Supplemental Figure 8, and data not shown). These findings are suggestive of a role for NRP2 in the regulation of effector CD4^+^ T cell activation.

**Figure 5:**
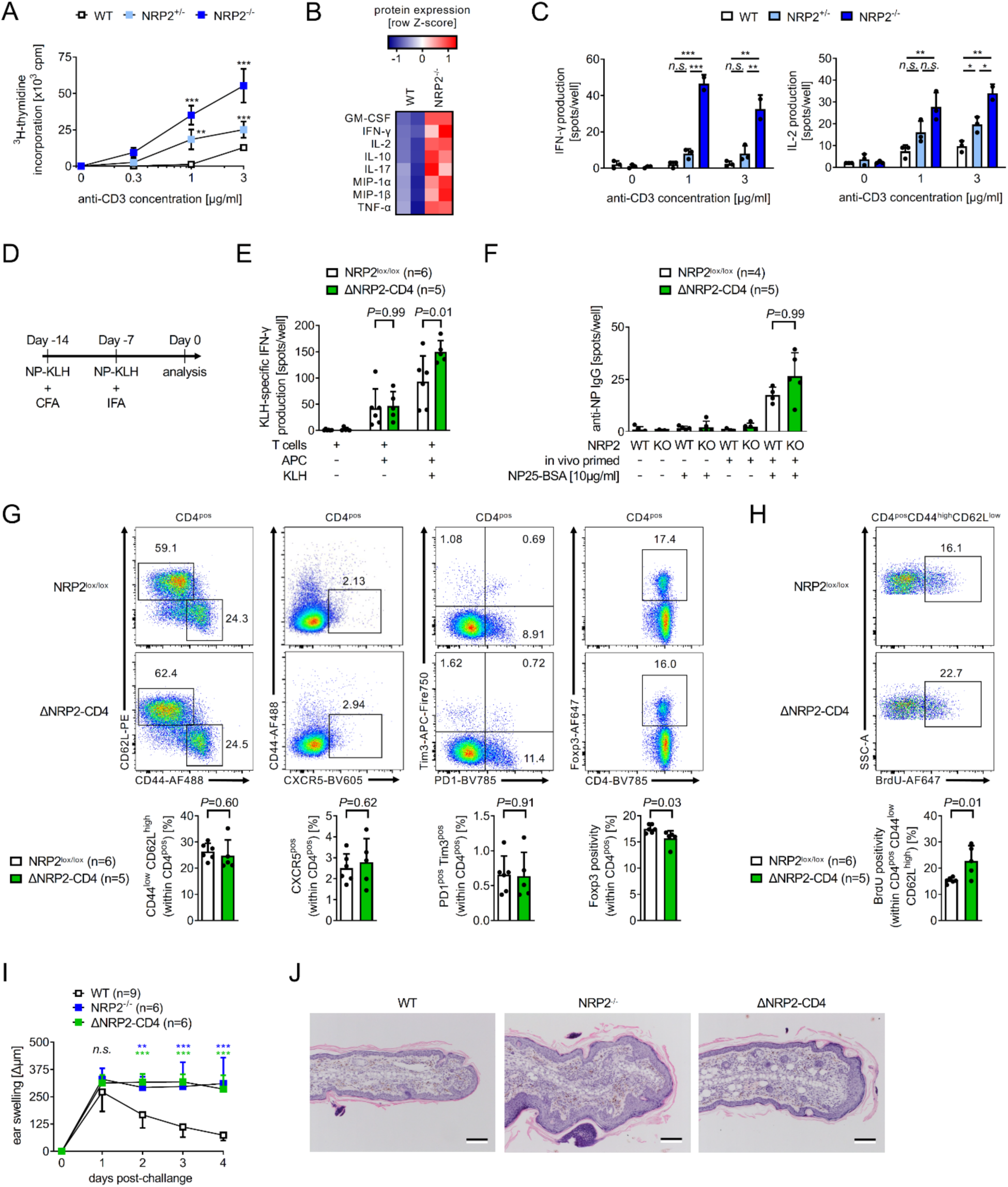
Knockout of NRP2 within CD4^+^ T cells increases pro-inflammatory responses in vitro and in vivo. **(A-C)** CD4^+^ T cells isolated from WT, heterozygous NRP2 knockout (NRP2^+/-^) and homozygous NRP2 knockout (NRP2^-/-^) mice were stimulated with increasing concentrations of plate-bound anti-CD3. **(A)** Proliferation as evaluated by ^3^H-thymidine incorporation after 72 hours (mean cpm ± SD from triplicate conditions; One-way ANOVA, ** *P*<0.01, *** *P*<0.001 vs. WT; representative of three independent experiments). **(B)** Cytokine concentrations in coculture supernatants from the experiments in Panel A (1 µg/ml anti-CD3) measured by multiplex-analyte profiling. The heatmap represents mean cytokine concentrations of duplicate conditions (two independent experiments). **(C)** IFN-γ and IL-2 production as assessed by ELISPOT (mean spots ± SD of triplicate condition; One-way ANOVA, *n.s.* not significant, ** *P*<0.01, *** *P*<0.001 vs. WT; representative of three independent experiments). **(D-H)** WT and ΔNRP2-CD4 knockout mice were immunized subcutaneously with NP-KLH (50 µg) in complete Freud’s adjuvant (CFA), boosted after 7 days with NP-KLH (50 µg) in incomplete Freud’s adjuvant (IFA), and T cell and B cell responses were analyzed after an additional 7 days. **(E)** IFN-γ production (by ELISPOT) following restimulation of primed CD4^+^ T cells to KLH. Bar graphs represent mean spots per well ± SD from NRP2^lox/lox^ (n=6) and ΔNRP2-CD4 (n=5) mice (Kruskal-Wallis test). **(F)** NP-specific IgG production in B cells by ELISPOT. Bar graphs represent mean spots per well ± SD from NRP2^lox/lox^ (n=4) and ΔNRP2-CD4 (n=5) mice (Kruskal-Wallis test). **(G)** Phenotype of splenic CD4^+^ T cell subsets. Representative dot plots (top panels) and bar graphs depicting differences between NRP2^lox/lox^ (n=6) and ΔNRP2-CD4 (n=5) mice (bottom panels; mean ± SD; unpaired t-test). **(H)** Proliferation (BrdU incorporation) of CD4^+^CD44^high^CD62L^low^ T effector/memory cells. Representative dot plots (top panels) and bar graphs depicting differences between NRP2^lox/lox^ (n=6) and ΔNRP2-CD4 (n=5) mice (bottom panels; mean ± SD; unpaired t-test). **(I-J**) WT, NRP2^-/-^ and ΔNRP2-CD4 knockout mice were sensitized to oxazolone and challenged by application to the right ear in a standard DTH model; vehicle application to the left ear served as a control. **(I)** Differences in thickness between the right (challenge) and left (control) ears were measured daily (Δµm; One-way ANOVA, *n.s.* not significant, ** *P*<0.01, *** *P*<0.001 vs. WT). **(J)** Hematoxylin and eosin (H&E) staining of challenged ears harvested on day 4 post-challenge (representative of n=3/condition).

We also evaluated antigen-specific responses in WT and ΔNRP2-CD4 KO mice following vaccination with Keyhole Limpet Hemocyanin (NP-KLH, Figure 5D-H). Seven days following a booster, KLH-specific CD4^+^ T cell IFN-γ responses were found to be significantly increased in ΔNRP2-CD4 KO mice vs. WT mice (Figure 5E; *P*=0.01). Anti-NP IgG B cell activity also trended higher in KO mice (Figure 5F), suggesting that NRP2 functions as a co-inhibitory molecule in vivo. There was no difference in the numbers of naïve CD44^low^CD62L^high^, CXCR5^+^ follicular (Tfh) or PD-1^+^Tim-3^+^ Texh or Foxp3^+^ Treg subsets following vaccination (Figure 5G). However, there was a significantly increased proliferation rate within CD44^high^CD62L^low^ T effector/memory cells in NRP2 KO mice compared to WT mice (Figure 5H). In vivo, antigen-specific delayed-type hypersensitivity (DTH) responses also demonstrated enhanced inflammation in both global NRP2^-/-^ KO and ΔNRP2-CD4 KO mice (*P*<0.001 vs. WT mice, Figure 5I-J). Ear swelling peaked 24- hours after rechallenge in WT control mice and subsided over a 3-day period (Figure 5I). In contrast, the DTH response/ear swelling failed to resolve by day 4 in KO mice, and edema and mononuclear infiltration persisted compared to WT controls (Figure 5J). Although it is possible that NRP2 may function on additional immune cell types, for example CD8^+^ T cells in our studies, these collective findings demonstrate that it functions to regulate antigen-specific effector CD4^+^ T cell activation and expansion in vitro, and an associated enhanced inflammatory response in vivo.

### Effector T cell responses in NRP2 knockout mice following transplantation

We next wished to evaluate the function of NRP2 in the regulation of alloimmunity, and whether it has biological effects in CD4^+^ Teff and/or Treg responses following transplantation. Initially, we transplanted B6.C-H-2^bm12^ hearts into C57BL/6 WT or NRP2 KO mice, as this single MHC class II mismatch results in a chronic insidious response that is dependent on the relative activity of both CD4^+^ Teff and Treg cells (40, 41). As illustrated in Figure 6A, we found that graft failure was accelerated when either global NRP2^-/-^ KO mice or ΔNRP2-CD4 KO mice were used as recipients (MST=34 days, *P*=0.01 and MST=21 days; *P*<0.001 respectively) vs. WT mice (MST >60 days). Also, there was a significant difference in graft survival (*P*=0.001) when ΔNRP2-CD4 KO mice were used as recipients vs. global NRP2^-/-^ KO mice, suggestive that its dominant regulatory effect in this model is related to its expression on CD4^+^ T cells (Figure 6A). Furthermore, inflammation was marked within allografts from ΔNRP2-CD4 KO recipients at early times post-transplant (Figure 6B), and the rejection response was associated with enhanced Teff priming as assessed by anti-donor IFN- γ production by recipient CD4^+^ T cells (*P*=0.002 vs. WT recipients, Figure 6C). To confirm that rejection in ΔNRP2-CD4 KO mice is not dependent on CD8^+^ T effectors, we treated ΔNRP2-CD4 KO recipients with anti-CD8 (both before and after transplantation) and find a similar survival/rejection pattern as non-depleted mice (Figure 6D-E, MST 17 days vs. >60 days in WT NRP2^lox/lox^ controls; *P*=0.001). These findings are suggestive that the expression of NRP2 on CD4^+^ Teff cells is required for long-term graft survival following transplantation.

**Figure 6:**
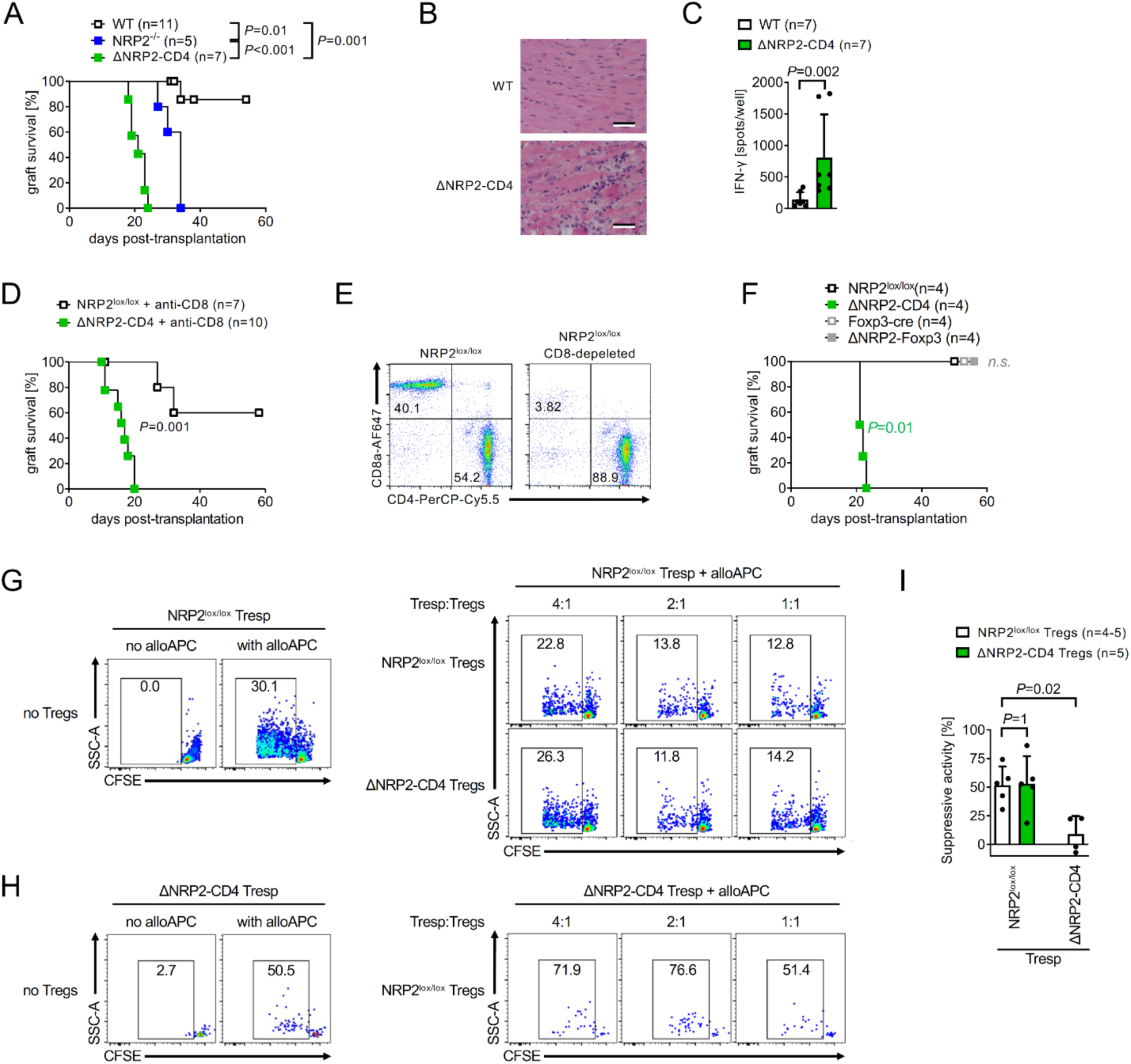
NRP2 expression on CD4^+^ T effector cells is required for long-term allograft survival. **(A-F)** Single MHC class II mismatched B6.C-H-2^bm12^ donor hearts were transplanted into NRP2 transgenic mice on the C57BL/6 background and graft function was monitored by palpation of the heartbeat. **(A)** Kaplan-Meier graft survival curves following transplantation of B6.C-H-2^bm12^ heart allografts into WT, NRP2^-/-^ or ΔNRP2-CD4 knockout recipients. **(B)** Histology as evaluated by H&E staining of allografts harvested on day 14 post-transplantation. **(C)** CD4^+^ T cell priming in WT and ΔNRP2-CD4 knockout mice as evaluated by IFN-γ ELISPOT on day 14 post-transplantation (mean spots/well ± SD; Mann Whitney test). **(D-E)** Allograft survival following transplantation of B6.C-H-2^bm12^ hearts into CD8-depleted ΔNRP2-CD4 knockout or NRP2^lox/lox^ recipients. **(E)** Efficiency of CD8 depletion from splenocytes by anti-CD8 treatment peri transplant by flow cytometry on day 2 post-transplantation. **(F)** Kaplan-Meier survival curves following transplantation of B6.C-H-2^bm12^ cardiac allografts into C57BL/6 NRP2^lox/lox^, ΔNRP2-CD4, Foxp3- cre or ΔNRP2-Foxp3 knockout recipients. **(G-I)** C57BL/6 NRP2^lox/lox^ and ΔNRP2-CD4 knockout mice received a fully MHC mismatched Balb/c skin transplant; Teff and Treg were harvested on day 14 and Treg function was assessed in an in vitro suppression assay. **(G-H)** A representative Treg suppression assay showing proliferation of NRP2 WT (G) or KO (H) Teff responders without Tregs (left panels) or with increasing ratios of Tregs (right panels). **(I)** Bar graph summarizing five independent assays comparing the % suppressive activity of NRP2^lox/lox^ and ΔNRP2-CD4 knockout Tregs (mean ± SD, One-way ANOVA with Tukey’s multiple comparison).

### NRP2 expression on Tregs is redundant for the modulation of alloimmunity

To determine if long-term graft survival is also dependent on NRP2-expressing CD4^+^ Treg subsets in vivo, we next performed transplants using ΔNRP2-Foxp3 KO mice as recipients of B6.C-H-2^bm12^ donor hearts. As illustrated in Figure 6F, we find that allografts survive long-term in ΔNRP2-Foxp3 KO recipients (MST >60 days, similar to WT recipients). Since long-term survival in this model is dependent on the expansion and function of Tregs (40, 41), this finding indicates that NRP2 is of major significance in the regulation of CD4^+^ Teff cell activity, and that its function on Tregs is redundant.

To further confirm that Tregs are functional in the absence of NRP2, we also performed in vitro suppression assays using alloprimed Teffs and pooled populations of CD25^high^ Tregs. Both cell types were isolated (day 14) from WT or KO C57BL/6 recipients of fully MHC mismatched Balb/c skin transplants, and proliferation of alloreactive CD4^+^ Teffs (+/- Tregs) was evaluated in a restimulation assay following coculture with allogeneic Balb/c APCs. In this manner, suppression by either NRP2 KO or WT Tregs can be compared, as previously reported (42). As illustrated in Figure 6G-I and Supplemental Figure 9, we find that both NRP2 KO and WT CD25^high^ Tregs are equally efficient in the suppression of WT effector CD4^+^ T responder cell proliferation. In contrast, we find ΔNRP2-CD4 KO Tresp cell proliferation is somewhat resistant to suppression by Tregs (Figure 6H-I). We also performed a validation experiment using FACS-sorted Foxp3-YFP^+^ Tregs instead of CD25^high^ Tregs. Identical to the findings in Figure 6G-I, we found that ΔNRP2-Foxp3 KO cells are functional to suppress WT Tresp, and that KO Tresp are resistant to Treg-mediated immunoregulation (Supplemental Figure 9C-F). These findings confirm that NRP2 deficient Treg cells are functional to suppress effector CD4^+^ T cells, and their suppressive potential is similar to WT Treg cells. Our findings also suggest that NRP2 deficient Teffs are intrinsically hyperactive.

Finally, in order to inhibit Teff priming, we treated ΔNRP2-CD4 KO or WT NRP2^lox/lox^ recipients of fully MHC mismatched Balb/c hearts with costimulatory blockade (anti-CD154 on days 0 and 2 post transplantation). We confirmed inhibition of allogeneic priming using a standard restimulation assay involving coculture with allogeneic Balb/c APCs (Supplemental Figure 10A- B). There was an almost identical graft survival pattern in KO and WT recipients following anti- CD154 treatment, suggestive that NRP2 is functional via its biology on alloactivated Teff subset(s). Similar to the B6.C-H-2^bm12^ model, graft survival following anti-CD154 treatment may also be dependent on Treg activity (43, 44). However, our observations demonstrate that Treg function is normal in the absence of NRP2, both in vivo (Figure 6F) and in vitro (Figure 6I), further indicating that the lack of difference in survival in this model is related to the inhibition of Teff priming/expansion.

### Independence of NRP2 and PD-1/PD-L1 co-inhibition

As discussed above, we find that NRP2 is co-expressed with known immune checkpoint/co-inhibitory molecules on phenotypically exhausted human and mouse CD4^+^ T cells. Since PD-1 is implicated in T cell dysfunction after transplantation (5), we also wished to determine if the regulatory effects of CD4^+^ T cell NRP2 is dependent on its association with PD-1. B6.C-H-2^bm12^ mice were used as donors and C57BL/6 WT or ΔNRP2-CD4 KO mice were used as recipients, either untreated or treated with anti-PD-L1. We found that anti-PD-L1 treatment of WT recipients resulted in accelerated rejection and early graft failure (MST = 17.5 days; *P*=0.02) vs. PBS-treated WT NRP2^lox/lox^ controls (MST >60 days; Figure 7). Anti-PD-L1 also reduced graft survival in ΔNRP2-CD4 KO recipients (MST=14 days) vs. in PBS-treated KO mice (MST=21 days; *P*=0.0002, Figure 7), and it resulted in a trend for accelerated rejection in ΔNRP2-CD4 KO mice vs. WT recipients (*P*=0.07, Figure 7). We interpret these observations to indicate that the co-inhibitory function of NRP2 is elicited through distinct signaling responses and/or regulatory ligands, independent of its association with PD-1.

**Figure 7:**
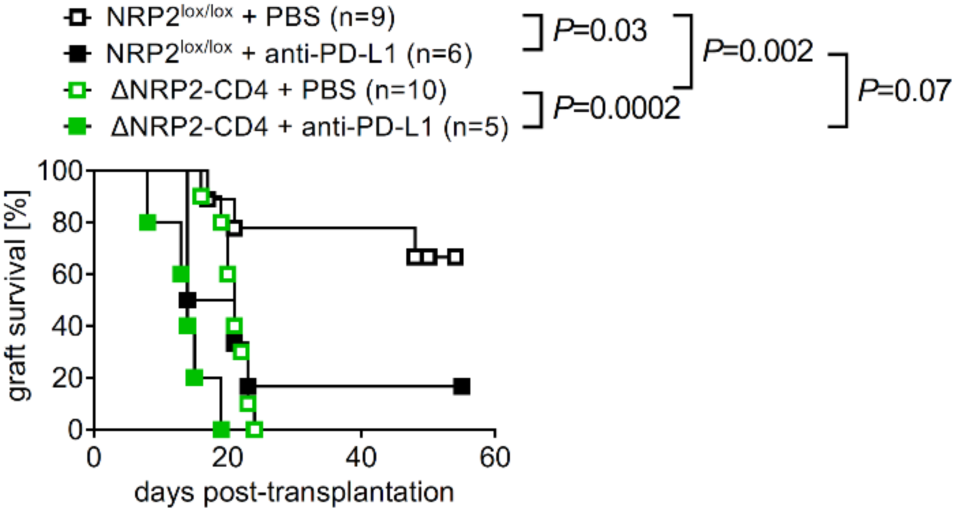
The regulation of allograft rejection by CD4^+^ T cell NRP2 is independent of PD- 1/PD-L1. Kaplan-Meier graft survival curves following the transplantation of B6.C-H-2^bm12^ heart allografts into ΔNRP2-CD4 or NRP2^lox/lox^ recipients that were treated with either anti-PD-L1 or PBS (on days 0, 3 and 6 post-transplantation).

## Discussion

Immunoregulatory mechanisms that promote long-term graft survival following solid organ transplantation are not fully understood. In this report, we find that NRP2 is expressed on a subset of human and mouse immune cell lineages, including T cells, B cells and APCs. We also find that it functions as a cell surface co-inhibitory receptor on CD4^+^ T cells to regulate cell-mediated and alloimmune responses. Furthermore, it is induced following vaccination on antigen-activated effector/memory CD4^+^ T cells, and its level of expression peaks on cells that are phenotypically and functionally exhausted. Knockout of NRP2 is associated with enhanced activation and proliferation in vitro, and enhanced inflammation in vivo in models of antigen-induced immunity and allograft rejection. Since NRP2 binds multiple ligands, these findings indicate that the relative expression of NRP2 on CD4^+^ T cell subsets has broad context-dependent implications for the regulation of inflammatory disease.

Our previous studies demonstrated that the NRP2 receptor elicits regulatory signaling responses in multiple cell types (including T cells) via the inhibition of PI3K/Akt/mTOR activity (45). Here, we find that knockout of NRP2 results in hyperactive CD4^+^ T cell responses in vitro, including enhanced IL-2 and IFN-γ production following mitogen activation and augmented T helper subset differentiation. These findings are consistent with the well-established function of PI3K/mTOR signaling in effector CD4^+^ cell responses (46–51). However, while activation of the mTOR signaling pathway inhibits Treg-dependent biology, we failed to observe any effect of NRP2 deficiency on the suppressive activity of Tregs in vitro or in vivo. Since NRP1 may also be expressed on Treg subsets (21, 33, 34, 36, 52) it is possible that NRP2-induced signaling is redundant inasmuch as both receptors elicit similar regulatory responses via common mechanisms (for example via the crosslinking of Plexin A family molecule(s) (18, 23, 24, 45, 53–56)). Thus, the loss of NRP2-dependent inhibition of Akt/mTOR signaling (45) may be compensated by ligand-dependent crosslinking of Plexin A or other molecules including NRP1 (53). Redundancy also explains why deletion of the NRP1 gene alone on Foxp3^+^ Treg cells does not result in autoimmune disease (34). Notably, in these studies, deletion of NRP2 did not result in any clinical signs of autoimmunity and phenotyping of knockout mice demonstrated normal T cell development (Supplemental Figure 7). Thus, the lack of autoimmunity in knockout mice is likely related to the redundancy of NRP2 on Tregs. Consistent with this interpretation, we note that this phenotype is similar to other coinhibitory molecule knockouts that do not develop autoimmunity but mount hyperactive antigen-specific immune responses (57). Although beyond the scope of the current studies, these observations are suggestive that the combined effects of NRP1 and NRP2 is necessary for Treg function which is critical for long-term transplant survival, the prevention of autoimmune disease and/or tumor immunity.

We also questioned why NRP2 is expressed by a small subset of CD4^+^ T cells at baseline, but deletion results in a generalized expansion following antigen-induced stimulation. One possibility is that NRP2 functions to limit the expansion of Teffs by promoting clonal deletion of Texh, an established mechanism that results in prolongation of graft survival (43). But deletion alone in the absence of immunoregulation is not necessarily sufficient to promote long-term graft survival (43). Thus, the expansion of Teffs in vivo in NRP2-deficient recipients may either be the result of a lack of Treg-dependent immunoregulation (40) and/or the deletion of Teffs as ascribed to other co-inhibitory molecules (58). However, as discussed above, we find that NRP2 knockout Tregs function as efficiently as wild type Tregs in the suppression of primed alloresponsive Teff cells. In contrast, NRP2-deficient Teff cells appear to be intrinsically hyperactive and are resistant to Treg-mediated suppression. Since NRP2 is induced on select populations of CD44^hi^ effector/memory CD4^+^ T cells, we interpret our findings to indicate that its main function is to augment clonal deletion of CD4^+^ Texh cells and/or to promote dysfunction. Consistent with this possibility, we also found that NRP2^pos^ cells are functionally characterized by reduced rates of proliferation and reduced cytokine production as compared to NRP2^neg^ T cells. Furthermore, they express a transcriptional program that includes multiple co-inhibitory receptors that augment exhaustion/dysfunction (4, 59, 60). Interestingly, it was reported that interferon regulatory factor 4 (IRF4) modulates CD4^+^ T cell differentiation into Texh (5, 61–64), and transcriptional data reveal that IRF4 KO CD4^+^ T cells express NRP2 (5). Indeed, the NRP2 promoter has 5 putative PU.1/IRF4 binding sites, and pilot analyses indicated that IRF4–NRP2 interactions are indeed inhibitory (data not shown). Thus, it is possible that when IRF4 is expressed (for example following early activation), it serves as a common mechanism to inhibit the expression of NRP2 and prevent the initiation of exhaustion. We thus suggest that the initial activation of CD4^+^ T cells results in IRF4-induced transcriptional activity that represses NRP2 expression; once IRF4 activity decreases, repression of NRP2 is removed allowing for ligand-dependent augmentation of T cell dysfunction and clonal deletion.

NRP2 ligands are broadly expressed within different tissues and may be functionally associated with the regulation of cell-mediated immune inflammation. For example, TGF-β is well established to be immunomodulatory (65), whereas VEGF-A and VEGF-C are generally associated with pro-inflammation (66, 67). Semaphorin family members are expressed within tumors where they are generally associated with immune evasion (39, 68–70). Pathologically, VEGF-A may bind NRP2 as an accessory receptor where it competitively occupies the semaphorin binding site and prevents semaphorin-dependent inhibitory signaling. In this manner, high levels of VEGF-A within a tissue (including allografts undergoing chronic rejection (67, 71, 72)) may inhibit SEMA3F–NRP2-induced immunomodulation. In contrast, high levels of semaphorins, for example within tumors, and/or soluble NRP2 that is present in the circulation (16, 73, 74) may competitively inhibit VEGF-VEGFR induced signaling and pro-inflammation (66). In our studies, we find a survival difference when either global NRP2 knockouts or our CD4^+^ T cell knockouts are used as recipients. We interpret this finding to indicate that NRP2 is also functional on other immune cell types (Figure 1A-B and Supplemental Figure 5B-D) and/or that soluble NRP2 functions as a scavenger receptor to inhibit rejection, for example by inhibiting pro-inflammatory VEGF-A (54, 66, 73). Nevertheless, since NRP2 ligands have potential to be expressed within allografts (72, 75), we performed an exploratory analysis of ligand expression using a dataset of human renal transplant biopsies (Supplemental Figure 11 (76, 77)). This analysis revealed that SEMA3F is among the highest expressed genes in stable/normal biopsies from patients on a tolerance induction protocol (Supplemental Figure 11B). These intriguing observations suggest that augmentation of the local expression of NRP2 ligands within allografts (including SEMA3F) may serve to sustain immunoregulation following transplantation.

In summary, NRP2 is expressed on a subset of antigen-activated effector/memory CD4^+^ T cells, and notably on CD4^+^ Texh cell subsets, where it functions to regulate alloimmunity and rejection following transplantation. It is also expressed on Treg cells, but its function is dominantly related to its biology on CD4^+^ Teff cells. We conclude that NRP2 signaling within CD4^+^ T cell subsets has implications for effector alloimmunity and thus long-term survival following transplantation.

## Methods

### Sex as a biological variable

Both males and females were used in all human and mouse studies, but donor-recipient sex mismatch was generally avoided in murine transplant studies.

### Isolation and cell culture of human peripheral blood mononuclear cells and CD4^+^ T cells

Human PBMC were isolated from healthy adult volunteer donors by high density centrifugation of blood (Corning, Corning, NY). CD4^+^ T cells were enriched and negatively isolated from human PBMCs using magnetic isolation beads (Stemcell Technologies, Vancouver, Canada) and purity of >98% was determined by flow cytometry. Cells were cultured in RPMI 1640 (Lonza, Walkersville, MD) supplemented with 10% FBS (Millipore Sigma, Darmstadt, Germany), 2 mM L-glutamine, 1 mM sodium pyruvate, 0.75 g/l sodium bicarbonate, 100 U/ml penicillin/streptomycin, 0.1 mM non-essential amino acids (all Lonza, Walkersville, MD) and 50 µM 2-mercaptoethanol (Millipore Sigma, Darmstadt, Germany) and activated using PHA (1-10 µg/ml; Remel, Thermo Scientific, Tewksbury, MA).

### Isolation and cell culture of murine CD4^+^ T cells

Murine CD4^+^ T cells were negatively isolated from splenocytes, as previously described (78). Cells were cultured in RPMI 1640 (Lonza, Walkersville, MD) supplemented with 10% FBS (Millipore Sigma, Darmstadt, Germany), 2 mM L-glutamine, 1 mM sodium pyruvate, 0.75 g/l sodium bicarbonate, 100 U/ml penicillin/streptomycin, 0.1 mM non-essential amino acids (all Lonza, Walkersville, MD) and 50 µM 2-mercaptoethanol (Millipore Sigma, Darmstadt, Germany). Murine T cells were stimulated with plate-bound anti-CD3 (clone 145-2C11; BioXcell, West Lebanon, NH) and proliferation was assessed by ^3^H-thymidine incorporation (1 µCi/well during last 16 hours of culture; scintillation counter: Wallac 1450 MicroBeta TriLux, version 4.6, Perkin Elmer, Boston, MA). Cytokine production was evaluated by either ELISPOT using the Ready-Set-Go Enzyme-linked immunospot assay kit (Thermo Scientific, Tewksbury, MA) according to manufacturer’s instructions, or by multiple analyte profiling using mouse cytokine/chemokine 25 plex magnetic bead panel (Millipore Sigma, Darmstadt, Germany) on a Luminex LX200 platform equipped with xPonent software (version 3.1.871.0; Luminex, Austin, TX). T helper cell polarization was performed as previously described (78). Briefly, naive CD4^+^ T cells were activated with plate-bound anti-CD3 and soluble anti-CD28 in polarizing conditions for 48 hours, rested in the absence of anti-CD3 for additional 48 hours, and Th differentiation was subsequently evaluated by stimulation with 10 µg/ml concanavalin A (Cayman Chemical Company, Ann Arbor, MI) in IFN- γ (Th1), IL-4 (Th2) or IL-17A (Th17) ELISPOT assays (all Thermo Scientific, Tewksbury, MA).

### Flow cytometry

Cells were washed twice in phosphate buffered saline (PBS) supplemented with 0.5% fetal bovine serum (FBS) and 6 µM EDTA (Boston BioProducts, Ashland, MA), preincubated with FcR blocker (BioLegend, San Diego, CA) for 20 minutes, stained with fluorochrome-conjugated antibodies (Supplemental Table 1) for 60 minutes at 4°C, washed, and analyzed within 24 hours. Specificity of anti-human NRP2 antibody staining was established using two different clones and in knock-down experiments (Supplemental Figure 12). Briefly, U87MG cells (ATCC, Manassas, VA) were cultured in EMEM supplemented with 10% FBS (Millipore Sigma, Darmstadt, Germany), and 100 U/ml penicillin/streptomycin (Lonza, Walkersville, MD) as described (45). NRP2 knockdown was performed with siRNA or scramble siRNA using lipofectamine in Opti-MEM (all Thermo Scientific, Tewksbuy, MA) for 24 hours; after washing, cells were cultured for an additional 3 days before analysis according to our previously reported protocol (45). The specificity of anti-murine NRP2 antibodies for use in cytometry were evaluated using murine NRP2 knockout cells (as detected by mRNA and Western blot analysis) and referenced to analysis using a NRP2-GFP-tag in transgenic mice. In general, we had concern that antibodies bind non-specifically to NRP2 knockout cells. Thus, for all flow-based studies, we used a single cell mRNA fluorescence in situ hybridization technique (PrimeFlow, Thermo Fisher Scientific, Tewksbury, MA) to evaluate NRP2 expression on immune cell subsets. PrimeFlow assays were performed using two independent probes targeting either exon 1-2 or exon 3-4 of the NRP2 transcript. Target specificity was confirmed using NRP2 KO cells (Supplemental Figure 6C). As a technical control, we used a probe targeting CD4 mRNA, and probe amplification in each sample was confirmed by comparing signals of CD4 mRNA *vs.* protein (not shown). We also used the GFP-tag in NRP2^lox/lox^ mice to identify NRP2 expression, but analysis of GFP was technically difficult due to the low signal/autofluorescence ratio of the GFP-tag. FoxP3 staining was performed using a FoxP3/transcription factor staining buffer set (Thermo Fisher, Tewksbury, MA). Intracellular cytokine staining was performed using an intracellular staining permeabilization buffer (BioLegend, San Diego, CA) on cells that were activated with 50 ng/ml phorbol 12-myristate 13-acetate and 1 μg/ml ionomycin (both MilliporeSigma, Burlington, MA) in the presence of 5 µg/ml brefeldin A (BioLegend, San Diego, CA) for 6 hours. BrdU staining was performed according to the manufacturer’s instructions using the phase-flow kit (BioLegend, San Diego, CA). Stained cells were analyzed either on a FACS Calibur (2 lasers), Celesta (3 lasers) or LSR II (5 lasers) cytometer (all BD Biosciences, San Diego, CA) and data were evaluated using FlowJo software (version v10.7, Tree Star, Ashland, OR). Cell sorting was performed on a FACS Aria II or Aria Fusion (both BD Biosciences, San Diego, CA) to a purity of >95%. Gating strategies and purity controls are depicted in each experiment.

### Immunocytology

For immunocytological analysis, CD4^+^ T cells were immobilized on ImmunoSelect adhesion slides (MoBiTec, Duesseldorf, Germany) for 20 minutes in PBS at 37°C. Adherent cells were subsequently fixed in 4% paraformaldehyde in PBS (Boston BioProducts, Ashland, MA) for 15 minutes at room temperature. After a wash in PBS, cells were permeabilized in cold methanol for 5 min at -20°C, washed and incubated with blocking buffer (5% BSA in PBS) for 30 minutes at room temperature and subsequently with primary antibodies (Supplemental Table 1) overnight at 4°C. After washing in PBS, cells were incubated with a species-specific fluorochrome-conjugated secondary antibody (Thermo Fisher, Tewksbury, MA) for 1 hour at room temperature, washed and mounted with ProLong Gold antifade containing 4’,6-diamidino- 2-phenylindole (Thermo Fisher, Tewksbury, MA). Staining was imaged on a confocal laser-scanning microscope (TCS SP5 X, Leica, Mannheim, Germany) equipped with LAS AF software (version 1.6.3; Leica, Mannheim, Germany). Specificity of NRP2 signal was evaluated using isotype control stainings.

### Western Blot analysis

Western blot analysis was performed using standard techniques and as previously described (78). Briefly, cells were lyzed in radioimmunoprecipitation assay buffer supplemented with EDTA and run on a 12% sodium dodecyl sulfate–polyacrylamide gel (Bio-Rad Laboratories, Hercules, CA). Proteins were transferred to a polyvinylidene difluoride membrane (MilliporeSigma, Burlington, MA), blocked with 10% BSA in TBS supplemented with 0.1% Tween 20 (TBST, Boston BioProducts, Milford, MA). Primary antibodies (see Supplemental Table 1) were diluted in blocking buffer prior to incubation overnight at 4°C. Following three washing steps in TBST, membranes were subsequently incubated with species-specific peroxidase-conjugated secondary antibodies (Jackson ImmunoResearch, West Grove, PA) for 1 hour at room temperature. After washing, the protein of interest was detected by chemiluminescence (Thermo Fisher, Tewksbury, MA) using a ChemiDoc MP imaging system (Bio-Rad, Hercules, CA). Membranes were stripped and reprobed to control for equal protein loading.

### Polymerase Chain Reaction (PCR)

PCR was performed using standard techniques. Briefly, DNA was extracted using the hot-alkaline DNA extraction method. Regions of interest were amplified using specific primer sets (Supplemental Table 2) and a master mix containing Taq DNA Polymerase, dNTPs and reaction buffer according to manufacturer’s instructions (Promega, Madison, WI). PCR products were run on an agarose gel containing SYBR green and visualized using a GelDoc XR+ imaging system (Bio-Rad, Hercules, CA).

### Transcriptomic analyses

For bulk RNA-sequencing, cells were sorted by flow cytometry staining, pelleted, lysed in TRIzol (Thermo Fisher, Tewksbury, MA) and stored at -80°C for group analysis. Library preparations using poly-A enriched RNA and sequencing was performed by Genewiz (South Plainfield, NJ). Paired-end 150bp reads were aligned to a reference human genome database (GRCh38) using the 2 pass STAR aligner (79) and gene expression levels were quantified using the GENCODE gene models (Release M14 (80)) using the HTseq count method (81). Normalization and differential gene expression was determined using the DESeq2 method (version 1.28.1 (82)) in the R Statistical Computing Environment (version 4.0.0). Functional enrichment, signaling pathway activities, and protein association network analyses were performed using the Gene set enrichment (83) and STRING (84) bioinformatics resources. One sample was identified as outlier using principal component analysis and subsequently removed from the analysis.

Single cell RNA-sequencing was combined with detection of NRP2 protein expression in a Cellular Indexing of Transcriptomes and Epitopes by Sequencing (CITE-seq) assay. Briefly, cells were preincubated with 10 µg/ml streptavidin and FcR blocker (both BioLegend, San Diego, CA) for 20 minutes, washed twice, stained with fluorochrome-conjugated antibodies for cell sorting, and PE- (clone MM03) and biotin- (clone 2v2) conjugated anti-NRP2 primary antibodies (Supplemental Table 1). Cells were washed and incubated with DNA-conjugated anti-PE and anti-biotin secondary antibodies (Supplemental Table 1). After washing, cells were FACS-sorted, washed again and resuspended in PBS supplemented with 1% bovine serum albumin (BSA). 3’- end mRNA and CITE-seq library preparations were immediately performed on a 10x Chromium platform (10x Genomics, Pleasanton, CA) and sequenced on a NextSeq 500 (Illumina, San Diego, CA). FASTQ files were demultiplexed and aligned to the human genome (GRCh38) using the Cell Ranger pipeline (version 3.1.0, 10x Genomics, Pleasanton, CA) and imported into Seurat (version 3.2.0; (85)) in the R Statistical Computing Environment (version 4.0.0). Cells with unusually high unique molecular identifiers (doublets), mitochondrial gene percentages (dying cells), and CD8^+^ T cells were excluded from further analysis. Predefined immune gene (86) expression was log-normalized, dimensional reduced, and clustered based on their k-nearest-neighbors. Markers for each cluster were ranked by their log fold change and cluster annotation was manually chosen based on gene expression profiles. Trajectory-based pseudotime analysis was performed using the Monocle3 method (version 0.2.2; (87)).

### Mouse strains

6-10-week-old C57BL/6 WT (CD45.2^+^) and congenic *Ptprc^a^Pepc^b^* (CD45.1^+^, both H-2b), Balb/c (H-2d), and B6.C-H-2^bm12^ mice were purchased from the Jackson Laboratory (Bar Harbor, ME). Ovalbumin-specific T cell receptor transgenic (OT-II) mice were gifted by Dr. Hans Oettgen (Boston Children’s Hospital, Boston, MA) and originally purchased from the Jackson Laboratory (Bar Harbor, ME). SCID-beige mice were purchased from Taconic (Germantown, NY). Transgenic NRP2 knock-out and NRP2 floxed mice were purchased from the Jackson Laboratory (Bar Harbor, ME) and backcrossed to >98% congenicity using C57BL/6 mice (384- SNP panel, Charles River, Wilmington, MA). NRP2 floxed mice were crossed with C57BL/6 CD4-cre (Taconic, Germantown, NY) and Foxp3-cre (Jackson Laboratory, Bar Harbor, ME) mice to generate conditional knock-out strains (ΔNRP2-CD4 and ΔNRP2-Foxp3 respectively). For each experiment, NRP2^lox/lox^ mice and 100% congenic C57BL/6 mice (Jackson Laboratory, Bar Harbor, ME) were included as controls.

### Humanized SCID-beige mice

Humanized SCID-beige mice (huSCID) were generated by the transfer of 5x10^7^ human peripheral blood mononuclear cells (PBMC) by tail vein injection. As a time course (generally up to 21 days) following humanization, mice were euthanized and the spleens were harvested, dissociated into a single cell solution, blocked with human and mouse FcR blocker and stained with fluorochrome conjugated antibodies as described above (see Supplemental Table 1). The phenotype of human CD4^+^ T cells within the spleen was assessed by gating on live human CD3^+^ CD4^+^ cells with gating out mouse CD45^+^ cells using flow cytometry. Alternatively, human CD4^+^ cells were FACS-sorted from splenocytes using the same gating strategy (human CD3^+^CD4^+^ cells/mouse CD45^neg^) for transcriptomic analysis.

### Heart transplantation

Heterotopic intraabdominal cardiac transplantation was performed as previously described (88) and graft survival was monitored by palpation of the heartbeat. In some recipients, CD8^+^ T cells were depleted by intraperitoneal injection of anti-CD8 (200 µg of clone 53-6.7; BioXcell, West Lebanon, NH) on day -5, -2 and +2 peri-transplant and every 4^th^ day thereafter. Efficacy of depletion was evaluated by flow cytometry. As indicated, some recipients were treated with anti-CD154 (200 µg/i.p; clone MR-1, BioXcell, West Lebanon, NH) on days 0 and 2 post-transplantation or anti-PD-L1 (200 µg/i.p; clone 10F.9G2, BioXcell, West Lebanon, NH) on day 0, 3 and 6 post-transplantation.

### Treg suppression assays

Allogeneic priming of T cells was performed by tail skin transplantion (as we described (78)) using Balb/c donors and C57BL/6 wild type or ΔNRP2-CD4 knockout mice as recipients. Spleens were harvested on day 14 post-transplantation and CD4^+^CD25^high^ Tregs, CD4^+^CD25^neg^ T responders (Tresp) were isolated as described above. Tresp were stained with CFSE (Thermo Fisher, Tewksbury, MA) prior to coculture with irradiated (1200 rad) CellTrace Far Red-stained (Thermo Fisher, Tewksbury, MA) Balb/c splenocytes. Tregs from WT or ΔNRP2- CD4 knockout recipients were added to cocultures in increasing ratios with Tresp (1:16 to 1:1) and suppression of Tresp proliferation was evaluated after 5 days by flow cytometry. For statistical purposes, suppressive activity was calculated as ([percentage of CFSE^dim^ cells in the absence of Tregs] - [percentage of CFSE^dim^ cells at 1:1 Tresp:Tregs])/[percentage of CFSE^dim^ cells in the absence of Tregs].

### Priming/Antigen-induced Responsiveness in vivo

Allogeneic priming of CD4^+^ T cells following cardiac transplantation was assessed by coculture of recipient CD4^+^ T cells (isolated from recipient splenocytes) in a mixed lymphocyte reaction with irradiated (1200 rad) donor splenocytes. Recipient CD4^+^ activity was either evaluated using CFSE dilution following five days of co-culture or by IFN-γ ELISPOT using anti-IFN-γ coated 96-well PVDF ELISPOT plates (Immobilon-P; MilliporeSigma, Burlington, MA). After 24 hours coculture, ELISPOT plates were washed twice with 0.1% Tween20 in PBS and subsequently incubated with a biotinylated anti-IFN-γ antibody (Supplemental Table 1). After 1 hour at 4°C, ELISPOT plates were washed twice with 0.1% Tween20 in PBS and subsequently incubated with HRP-conjugated avidin for 45 minutes. Following two washing steps with 0.1% Tween20 in PBS and then PBS, HRP-activity was developed using 3-Amino-9-ethylcarbazole (AEC, BD Bioscience, San Jose, CA). Stained plates were scanned and analyzed on an ImmunoSpot S6 Ultra ELISPOT reader (version 5.0, CTL, Shaker Heights, OH).

In some experiments, ovalbumin-specific T cell receptor transgenic OT-II CD4^+^ T cells (2.5x10^6^ cells/mouse) were adoptively transferred by tail vein injection into congenic C57BL/6 Ptprc^a^Pepc^b^ (CD45.1^pos^CD45.2^neg^) host mice. After 24hrs, host mice were immunized with 50 μg ovalbumin in Complete Freund’s Adjuvant (CFA, InvivoGen, San Diego, CA) subcutaneously in the right flank. Subsequently, NRP2 expression and the phenotype of antigen-specific (identified using anti-CD45.2) vs. host (identified using anti-CD45.1) CD4^+^ T cells was evaluated in splenocytes as a time course up to 7 days post-vaccination.

Antigen-induced responsiveness was assessed following vaccination, where mice were immunized with 50 μg NP-KLH (NP:KLH ratio: 24:1; LGC Biosearch Technologies, Hoddesdon, United Kingdom) emulsified in CFA subcutaneously in the right flank. After 7 days, the mice received a booster of NP-KLH (50 μg) emulsified in Incomplete Freund’s Adjuvant (InvivoGen, San Diego, CA). Following an additional 7 days, the mice were euthanized, and CD4^+^ T cells and CD19^+^ B cells were purified from the spleen by negative selection using magnetic bead isolation kits (Stemcell Technologies, Vancouver, Canada). In vivo priming to KLH was assessed by in vitro stimulation of CD4^+^ T cells with KLH for 24 hours and IFN-γ production was assessed by ELISPOT, as described above. B cells were stimulated in NP-conjugated bovine serum albumin (NP-BSA, 1 µg/ml; ratio: 25:1; LGC Biosearch Technologies, Hoddesdon, United Kingdom) in coated 96-well PVDF plates for 24 hours, and responses were assessed by ELISPOT following incubation with 4 μg/ml HRP-conjugated anti-mouse IgG for 1 hour at 4°C (Supplemental Table 1). As above, cells were washed both before and after secondary antibody incubation and prior to development using AEC (BD Bioscience, San Jose, CA). In some experiments, 1 mg BrdU (BioLegend, San Diego, CA) was injected intraperitoneally every 12 hours for the last 72 hours and proliferation as well as T cell subset phenotype were analyzed using flow cytometry.

### Delayed-type hypersensitivity

Delayed-type hypersensitivity (DTH) responses were performed using an established model (89). Briefly mice were sensitized using 2% 4-ethoxymethylene-2- phenyl-2-oxazolin-5-one (oxazolone; dissolved in 20% olive oil in acetone; Millipore Sigma, Darmstadt, Germany) applied to the shaved abdomen (50 µl) and each paw (5 µl). Five days after sensitization, DTH responses were evaluated in the right ear following reapplication of a 1% oxazolone solution (10 µl); the left ear was treated with vehicle alone as a control. The thicknesses of the ears were measured daily with a caliper (dial thickness gauge; Swiss Precision Instruments, Garden Grove, CA) and are expressed as difference between oxazolone-challenged vs. vehicle control (Δµm).

### Histology

Tissue was harvested and fixed in 10% formaldehyde in PBS (Fisher Scientific, Kalamazoo, MI) overnight at 4°C, paraffin-embedded, sectioned and stained with Hematoxylin and Eosin (all Thermo Fisher, Tewksbury, MA). Staining was imaged on a Nikon Eclipse 80i (Nikon, Mississauga, Ontario, Canada) using a Retiga-2000R CCD camera (QImaging, Surrey, Canada) equipped with NIS Elements software (version 3.22.15; Nikon, Melville, NY).

### Statistics

Statistical analyses were performed using the two-tailed Student’s t-test or One-way ANOVA as indicated with previous testing of equality of variances. The Mann-Whitney test or Kruskal-Wallis test were used if variances were significantly different. Graft survival was analyzed using the Log-Rank test. *P*-values of less than 0.05 were considered significant. Heatmaps were generated using the heatmap.2 function in the gplots package (version 3.0.4) using the R Statistical Computing Environment (version 4.0.0).

## Study approval

Volunteer blood donors were consented in accordance with Institutional Review Board approval at Boston Children’s Hospital. All animal studies outlined in this publication using mice were approved by the Institutional Animal Care and Use Committee at Boston Children’s Hospital and comply with the NIH Guide for the Care and Use of Laboratory Animals.

## Data Availability Statement

Raw datasets generated in this study are available in the National Center for Biotechnology Information (NCBI) Gene Expression Omnibus (GEO) repository https://www.ncbi.nlm.nih.gov/geo/query/acc.cgi?acc=GSE231735 under the accession no. GSE231735. Transcriptomic data from renal graft biopsies were reanalyzed from publicly available data (GSE106675). The graphical abstract was created with BioRender.com.

## Author contributions

JW and DMB conceptualized the study and designed the experiments. JW, NK, SB, JL, MM, BA, KM, KL, HZ, MK, and HN conducted experiments, and JW, NK, MM, BA, and KM acquired data. JW, NK, SWK, DMB analyzed data. DRB provided reagents and mice, supported analysis of data. JW and DMB wrote and edited the manuscript.

## Acknowledgements

This work was supported by the National Institutes of Health (R01AI148539), the Isabella Julian Forrest Foundation and the Casey Lee Ball Foundation. The authors wish to thank Dr. Dakshnapriya Balasubbramanian and Dr. In-Hee Lee (both Boston Children’s Hospital, Boston, MA) for assistance with data analysis, Dr. Andrea Cubitt (aTyr Pharma, San Diego, CA) for gifting an anti-human NRP2 antibody (clone 2v2) and Drs. Hans Oettgen and Cynthia Kanagaratham (both Boston Children’s Hospital, Boston, MA) for gifting OT-II transgenic mice. We also wish to acknowledge Evan Chernov, Megan Cooper, Evelyn Flynn, Dr. Jessica Leyva-Rangel, Dr. Yao Gao, Mahima Poreddy and Dr. Liang Zhou (all Boston Children’s Hospital, Boston, MA) for technical support for both in vitro and in vivo studies.

## References

1. Blank CU, Haining WN, Held W, Hogan PG, Kallies A, Lugli E, et al. Defining ’T cell exhaustion’. Nature reviews Immunology. 2019;19(11):665–74.

2. Crawford A, Angelosanto JM, Kao C, Doering TA, Odorizzi PM, Barnett BE, et al. Molecular and transcriptional basis of CD4(+) T cell dysfunction during chronic infection. Immunity. 2014;40(2):289–302.

3. Chihara N, Madi A, Kondo T, Zhang H, Acharya N, Singer M, et al. Induction and transcriptional regulation of the co-inhibitory gene module in T cells. Nature. 2018;558(7710):454-9.

4. Tian X, Zhang A, Qiu C, Wang W, Yang Y, Qiu C, et al. The upregulation of LAG-3 on T cells defines a subpopulation with functional exhaustion and correlates with disease progression in HIV-infected subjects. J Immunol. 2015;194(8):3873–82.

5. Wu J, Zhang H, Shi X, Xiao X, Fan Y, Minze LJ, et al. Ablation of Transcription Factor IRF4 Promotes Transplant Acceptance by Driving Allogenic CD4(+) T Cell Dysfunction. Immunity. 2017;47(6):1114–28 e6.

6. McKinney EF, Lee JC, Jayne DR, Lyons PA, and Smith KG. T-cell exhaustion, co-stimulation and clinical outcome in autoimmunity and infection. Nature. 2015;523(7562):612-6.

7. Rangachari M, Zhu C, Sakuishi K, Xiao S, Karman J, Chen A, et al. Bat3 promotes T cell responses and autoimmunity by repressing Tim-3-mediated cell death and exhaustion. Nature medicine. 2012;18(9):1394–400.

8. Zajac AJ, Blattman JN, Murali-Krishna K, Sourdive DJ, Suresh M, Altman JD, et al. Viral immune evasion due to persistence of activated T cells without effector function. J Exp Med. 1998;188(12):2205–13.

9. Moskophidis D, Lechner F, Pircher H, and Zinkernagel RM. Virus persistence in acutely infected immunocompetent mice by exhaustion of antiviral cytotoxic effector T cells. Nature. 1993;362(6422):758-61.

10. Willimsky G, and Blankenstein T. Sporadic immunogenic tumours avoid destruction by inducing T-cell tolerance. Nature. 2005;437(7055):141-6.

11. Fourcade J, Sun Z, Benallaoua M, Guillaume P, Luescher IF, Sander C, et al. Upregulation of Tim-3 and PD-1 expression is associated with tumor antigen-specific CD8+ T cell dysfunction in melanoma patients. J Exp Med. 2010;207(10):2175–86.

12. Fribourg M, Anderson L, Fischman C, Cantarelli C, Perin L, La Manna G, et al. T-cell exhaustion correlates with improved outcomes in kidney transplant recipients. Kidney Int. 2019;96(2):436–49.

13. Steger U, Denecke C, Sawitzki B, Karim M, Jones ND, and Wood KJ. Exhaustive differentiation of alloreactive CD8+ T cells: critical for determination of graft acceptance or rejection. Transplantation. 2008;85(9):1339–47.

14. Sarraj B, Ye J, Akl AI, Chen G, Wang JJ, Zhang Z, et al. Impaired selectin-dependent leukocyte recruitment induces T-cell exhaustion and prevents chronic allograft vasculopathy and rejection. Proceedings of the National Academy of Sciences of the United States of America. 2014;111(33):12145–50.

15. Miller ML, McIntosh CM, Wang Y, Chen L, Wang P, Lei YM, et al. Resilience of T cell-intrinsic dysfunction in transplantation tolerance. Proceedings of the National Academy of Sciences of the United States of America. 2019;116(47):23682–90.

16. Rossignol M, Gagnon ML, and Klagsbrun M. Genomic organization of human neuropilin- 1 and neuropilin-2 genes: identification and distribution of splice variants and soluble isoforms. Genomics. 2000;70(2):211–22.

17. He Z, and Tessier-Lavigne M. Neuropilin is a receptor for the axonal chemorepellent Semaphorin III. Cell. 1997;90(4):739–51.

18. Kolodkin AL, Levengood DV, Rowe EG, Tai YT, Giger RJ, and Ginty DD. Neuropilin is a semaphorin III receptor. Cell. 1997;90(4):753–62.

19. Chen H, Chedotal A, He Z, Goodman CS, and Tessier-Lavigne M. Neuropilin-2, a novel member of the neuropilin family, is a high affinity receptor for the semaphorins Sema E and Sema IV but not Sema III. Neuron. 1997;19(3):547–59.

20. Soker S, Takashima S, Miao HQ, Neufeld G, and Klagsbrun M. Neuropilin-1 is expressed by endothelial and tumor cells as an isoform-specific receptor for vascular endothelial growth factor. Cell. 1998;92(6):735–45.

21. Glinka Y, and Prud’homme GJ. Neuropilin-1 is a receptor for transforming growth factor beta-1, activates its latent form, and promotes regulatory T cell activity. J Leukoc Biol. 2008;84(1):302–10.

22. Guo HF, and Vander Kooi CW. Neuropilin Functions as an Essential Cell Surface Receptor. The Journal of biological chemistry. 2015;290(49):29120–6.

23. Parker MW, Xu P, Li X, and Vander Kooi CW. Structural basis for selective vascular endothelial growth factor-A (VEGF-A) binding to neuropilin-1. The Journal of biological chemistry. 2012;287(14):11082–9.

24. Janssen BJ, Malinauskas T, Weir GA, Cader MZ, Siebold C, and Jones EY. Neuropilins lock secreted semaphorins onto plexins in a ternary signaling complex. Nat Struct Mol Biol. 2012;19(12):1293–9.

25. Bielenberg DR, Pettaway CA, Takashima S, and Klagsbrun M. Neuropilins in neoplasms: expression, regulation, and function. Exp Cell Res. 2006;312(5):584–93.

26. Matthies AM, Low QE, Lingen MW, and DiPietro LA. Neuropilin-1 participates in wound angiogenesis. The American journal of pathology. 2002;160(1):289–96.

27. Wang L, Zeng H, Wang P, Soker S, and Mukhopadhyay D. Neuropilin-1-mediated vascular permeability factor/vascular endothelial growth factor-dependent endothelial cell migration. The Journal of biological chemistry. 2003;278(49):48848–60.

28. Bachelder RE, Lipscomb EA, Lin X, Wendt MA, Chadborn NH, Eickholt BJ, et al. Competing autocrine pathways involving alternative neuropilin-1 ligands regulate chemotaxis of carcinoma cells. Cancer Res. 2003;63(17):5230–3.

29. Parikh AA, Fan F, Liu WB, Ahmad SA, Stoeltzing O, Reinmuth N, et al. Neuropilin-1 in human colon cancer: expression, regulation, and role in induction of angiogenesis. The American journal of pathology. 2004;164(6):2139–51.

30. Kumanogoh A, and Kikutani H. Immunological functions of the neuropilins and plexins as receptors for semaphorins. Nature reviews Immunology. 2013;13(11):802–14.

31. Roy S, Bag AK, Singh RK, Talmadge JE, Batra SK, and Datta K. Multifaceted Role of Neuropilins in the Immune System: Potential Targets for Immunotherapy. Frontiers in immunology. 2017;8:1228.

32. Hill JA, Feuerer M, Tash K, Haxhinasto S, Perez J, Melamed R, et al. Foxp3 transcription-factor-dependent and -independent regulation of the regulatory T cell transcriptional signature. Immunity. 2007;27(5):786–800.

33. Sarris M, Andersen KG, Randow F, Mayr L, and Betz AG. Neuropilin-1 expression on regulatory T cells enhances their interactions with dendritic cells during antigen recognition. Immunity. 2008;28(3):402–13.

34. Delgoffe GM, Woo SR, Turnis ME, Gravano DM, Guy C, Overacre AE, et al. Stability and function of regulatory T cells is maintained by a neuropilin-1-semaphorin-4a axis. Nature. 2013;501(7466):252-6.

35. Overacre-Delgoffe AE, Chikina M, Dadey RE, Yano H, Brunazzi EA, Shayan G, et al. Interferon-gamma Drives Treg Fragility to Promote Anti-tumor Immunity. Cell. 2017;169(6):1130–41 e11.

36. Yadav M, Louvet C, Davini D, Gardner JM, Martinez-Llordella M, Bailey-Bucktrout S, et al. Neuropilin-1 distinguishes natural and inducible regulatory T cells among regulatory T cell subsets in vivo. J Exp Med. 2012;209(10):1713–22, S1-19.

37. Solomon BD, Mueller C, Chae WJ, Alabanza LM, and Bynoe MS. Neuropilin-1 attenuates autoreactivity in experimental autoimmune encephalomyelitis. Proceedings of the National Academy of Sciences of the United States of America. 2011;108(5):2040–5.

38. Liu C, Somasundaram A, Manne S, Gocher AM, Szymczak-Workman AL, Vignali KM, et al. Neuropilin-1 is a T cell memory checkpoint limiting long-term antitumor immunity. Nature immunology. 2020;21(9):1010–21.

39. Barnkob MB, Michaels YS, Andre V, Macklin PS, Gileadi U, Valvo S, et al. Semmaphorin 3 A causes immune suppression by inducing cytoskeletal paralysis in tumour-specific CD8(+) T cells. Nat Commun. 2024;15(1):3173.

40. Schenk S, Kish DD, He C, El-Sawy T, Chiffoleau E, Chen C, et al. Alloreactive T cell responses and acute rejection of single class II MHC-disparate heart allografts are under strict regulation by CD4+ CD25+ T cells. J Immunol. 2005;174(6):3741–8.

41. 41. Wedel J, Stack MP, Seto T, Sheehan MM, Flynn EA, Stillman IE, et al. T Cell-Specific Adaptor Protein Regulates Mitochondrial Function and CD4(+) T Regulatory Cell Activity In Vivo following Transplantation. J Immunol. 2019;203(8):2328–38.

42. Collison LW, and Vignali DA. In vitro Treg suppression assays. Methods in molecular biology. 2011;707:21–37.

43. Quezada SA, Bennett K, Blazar BR, Rudensky AY, Sakaguchi S, and Noelle RJ. Analysis of the underlying cellular mechanisms of anti-CD154-induced graft tolerance: the interplay of clonal anergy and immune regulation. J Immunol. 2005;175(2):771–9.

44. Taylor PA, Friedman TM, Korngold R, Noelle RJ, and Blazar BR. Tolerance induction of alloreactive T cells via ex vivo blockade of the CD40:CD40L costimulatory pathway results in the generation of a potent immune regulatory cell. Blood. 2002;99(12):4601–9.

45. Nakayama H, Bruneau S, Kochupurakkal N, Coma S, Briscoe DM, and Klagsbrun M. Regulation of mTOR Signaling by Semaphorin 3F-Neuropilin 2 Interactions In Vitro and In Vivo. Scientific reports. 2015;5:11789.

46. Delgoffe GM, Kole TP, Zheng Y, Zarek PE, Matthews KL, Xiao B, et al. The mTOR kinase differentially regulates effector and regulatory T cell lineage commitment. Immunity. 2009;30(6):832–44.

47. Delgoffe GM, Pollizzi KN, Waickman AT, Heikamp E, Meyers DJ, Horton MR, et al. The kinase mTOR regulates the differentiation of helper T cells through the selective activation of signaling by mTORC1 and mTORC2. Nature immunology. 2011;12(4):295–303.

48. Delgoffe GM, and Powell JD. Exploring functional in vivo consequences of the selective genetic ablation of mTOR signaling in T helper lymphocytes. Methods in molecular biology. 2012;821:317–27.

49. Zeng H, Yang K, Cloer C, Neale G, Vogel P, and Chi H. mTORC1 couples immune signals and metabolic programming to establish T(reg)-cell function. Nature. 2013;499(7459):485-90.

50. Shrestha S, Yang K, Guy C, Vogel P, Neale G, and Chi H. Treg cells require the phosphatase PTEN to restrain TH1 and TFH cell responses. Nature immunology. 2015;16(2):178–87.

51. Huynh A, DuPage M, Priyadharshini B, Sage PT, Quiros J, Borges CM, et al. Control of PI(3) kinase in Treg cells maintains homeostasis and lineage stability. Nature immunology. 2015;16(2):188–96.

52. Hansen W, Hutzler M, Abel S, Alter C, Stockmann C, Kliche S, et al. Neuropilin 1 deficiency on CD4+Foxp3+ regulatory T cells impairs mouse melanoma growth. J Exp Med. 2012;209(11):2001–16.

53. Antipenko A, Himanen JP, van Leyen K, Nardi-Dei V, Lesniak J, Barton WA, et al. Structure of the semaphorin-3A receptor binding module. Neuron. 2003;39(4):589–98.

54. Parker MW, Linkugel AD, Goel HL, Wu T, Mercurio AM, and Vander Kooi CW. Structural basis for VEGF-C binding to neuropilin-2 and sequestration by a soluble splice form. Structure. 2015;23(4):677–87.

55. Sulpice E, Plouet J, Berge M, Allanic D, Tobelem G, and Merkulova-Rainon T. Neuropilin-1 and neuropilin-2 act as coreceptors, potentiating proangiogenic activity. Blood. 2008;111(4):2036–45.

56. Yamamoto M, Suzuki K, Okuno T, Ogata T, Takegahara N, Takamatsu H, et al. Plexin- A4 negatively regulates T lymphocyte responses. International immunology. 2008;20(3):413–20.

57. Keir ME, Freeman GJ, and Sharpe AH. PD-1 regulates self-reactive CD8+ T cell responses to antigen in lymph nodes and tissues. J Immunol. 2007;179(8):5064–70.

58. Tai X, Van Laethem F, Pobezinsky L, Guinter T, Sharrow SO, Adams A, et al. Basis of CTLA-4 function in regulatory and conventional CD4(+) T cells. Blood. 2012;119(22):5155–63.

59. Wolf Y, Anderson AC, and Kuchroo VK. TIM3 comes of age as an inhibitory receptor. Nature reviews Immunology. 2020;20(3):173–85.

60. Sharpe AH, and Pauken KE. The diverse functions of the PD1 inhibitory pathway. Nature reviews Immunology. 2018;18(3):153–67.

61. Brustle A, Heink S, Huber M, Rosenplanter C, Stadelmann C, Yu P, et al. The development of inflammatory T(H)-17 cells requires interferon-regulatory factor 4. Nature immunology. 2007;8(9):958–66.

62. Grusdat M, McIlwain DR, Xu HC, Pozdeev VI, Knievel J, Crome SQ, et al. IRF4 and BATF are critical for CD8(+) T-cell function following infection with LCMV. Cell death and differentiation. 2014;21(7):1050–60.

63. Mittrucker HW, Matsuyama T, Grossman A, Kundig TM, Potter J, Shahinian A, et al. Requirement for the transcription factor LSIRF/IRF4 for mature B and T lymphocyte function. Science. 1997;275(5299):540-3.

64. Staudt V, Bothur E, Klein M, Lingnau K, Reuter S, Grebe N, et al. Interferon-regulatory factor 4 is essential for the developmental program of T helper 9 cells. Immunity. 2010;33(2):192–202.

65. Rubtsov YP, and Rudensky AY. TGFbeta signalling in control of T-cell-mediated self-reactivity. Nature reviews Immunology. 2007;7(6):443–53.

66. Reinders ME, Sho M, Izawa A, Wang P, Mukhopadhyay D, Koss KE, et al. Proinflammatory functions of vascular endothelial growth factor in alloimmunity. The Journal of clinical investigation. 2003;112(11):1655–65.

67. Bruneau S, Woda CB, Daly KP, Boneschansker L, Jain NG, Kochupurakkal N, et al. Key Features of the Intragraft Microenvironment that Determine Long-Term Survival Following Transplantation. Frontiers in immunology. 2012;3:54.

68. Casazza A, Laoui D, Wenes M, Rizzolio S, Bassani N, Mambretti M, et al. Impeding macrophage entry into hypoxic tumor areas by Sema3A/Nrp1 signaling blockade inhibits angiogenesis and restores antitumor immunity. Cancer cell. 2013;24(6):695–709.

69. Sierra JR, Corso S, Caione L, Cepero V, Conrotto P, Cignetti A, et al. Tumor angiogenesis and progression are enhanced by Sema4D produced by tumor-associated macrophages. J Exp Med. 2008;205(7):1673–85.

70. Wallerius M, Wallmann T, Bartish M, Ostling J, Mezheyeuski A, Tobin NP, et al. Guidance Molecule SEMA3A Restricts Tumor Growth by Differentially Regulating the Proliferation of Tumor-Associated Macrophages. Cancer Res. 2016;76(11):3166–78.

71. Daly KP, Stack M, Eisenga MF, Keane JF, Zurakowski D, Blume ED, et al. Vascular endothelial growth factor A is associated with the subsequent development of moderate or severe cardiac allograft vasculopathy in pediatric heart transplant recipients. The Journal of heart and lung transplantation : the official publication of the International Society for Heart Transplantation. 2017;36(4):434–42.

72. Wedel J, Bruneau S, Kochupurakkal N, Boneschansker L, and Briscoe DM. Chronic allograft rejection: a fresh look. Current opinion in organ transplantation. 2015;20(1):13–20.

73. Geretti E, van Meeteren LA, Shimizu A, Dudley AC, Claesson-Welsh L, and Klagsbrun M. A mutated soluble neuropilin-2 B domain antagonizes vascular endothelial growth factor bioactivity and inhibits tumor progression. Mol Cancer Res. 2010;8(8):1063–73.

74. Mayr T, Dix P, Hasheminasab S-L, Hampel C, Sylvester M, Schneberger N, et al. Discovery of a novel soluble Neuropilin-2 isoform with antiangiogenic and antitumorigenic activity. Presented at the Proceedings of the American Association for Cancer Research Annual Meeting; April 8-13, 2022; Philadelphia, Pennsylvania, USA. Cancer Res. 2022;82(12_Supplement):831.

75. Wedel J, Nakayama H, Kochupurakkal NM, Koch J, Klagsbrun M, Bielenberg DR, et al. The intragraft microenvironment as a central determinant of chronic rejection or local immunoregulation/tolerance. Current opinion in organ transplantation. 2017;22(1):55–63.

76. Gallon L, Mathew JM, Bontha SV, Dumur CI, Dalal P, Nadimpalli L, et al. Intragraft Molecular Pathways Associated with Tolerance Induction in Renal Transplantation. J Am Soc Nephrol. 2018;29(2):423–33.

77. Leventhal J, Abecassis M, Miller J, Gallon L, Ravindra K, Tollerud DJ, et al. Chimerism and tolerance without GVHD or engraftment syndrome in HLA-mismatched combined kidney and hematopoietic stem cell transplantation. Sci Transl Med. 2012;4(124):124ra28.

78. Wedel J, Bruneau S, Liu K, Kong SW, Sage PT, Sabatini DM, et al. DEPTOR modulates activation responses in CD4(+) T cells and enhances immunoregulation following transplantation. American journal of transplantation : official journal of the American Society of Transplantation and the American Society of Transplant Surgeons. 2019;19(1):77–88.

79. Dobin A, Davis CA, Schlesinger F, Drenkow J, Zaleski C, Jha S, et al. STAR: ultrafast universal RNA-seq aligner. Bioinformatics. 2013;29(1):15–21.

80. Mudge JM, and Harrow J. Creating reference gene annotation for the mouse C57BL6/J genome assembly. Mamm Genome. 2015;26(9-10):366–78.

81. Anders S, Pyl PT, and Huber W. HTSeq--a Python framework to work with high-throughput sequencing data. Bioinformatics. 2015;31(2):166–9.

82. Love MI, Huber W, and Anders S. Moderated estimation of fold change and dispersion for RNA-seq data with DESeq2. Genome Biol. 2014;15(12):550.

83. Subramanian A, Tamayo P, Mootha VK, Mukherjee S, Ebert BL, Gillette MA, et al. Gene set enrichment analysis: a knowledge-based approach for interpreting genome-wide expression profiles. Proceedings of the National Academy of Sciences of the United States of America. 2005;102(43):15545–50.

84. Szklarczyk D, Franceschini A, Wyder S, Forslund K, Heller D, Huerta-Cepas J, et al. STRING v10: protein-protein interaction networks, integrated over the tree of life. Nucleic Acids Res. 2015;43(D1):D447–52.

85. Stuart T, Butler A, Hoffman P, Hafemeister C, Papalexi E, Mauck WM, 3rd, et al. Comprehensive Integration of Single-Cell Data. Cell. 2019;177(7):1888–902 e21.

86. Miao YR, Zhang Q, Lei Q, Luo M, Xie GY, Wang H, et al. ImmuCellAI: A Unique Method for Comprehensive T-Cell Subsets Abundance Prediction and its Application in Cancer Immunotherapy. Adv Sci (Weinh). 2020;7(7):1902880.

87. Cao J, Spielmann M, Qiu X, Huang X, Ibrahim DM, Hill AJ, et al. The single-cell transcriptional landscape of mammalian organogenesis. Nature. 2019;566(7745):496–502.

88. Corry RJ, Winn HJ, and Russell PS. Primarily vascularized allografts of hearts in mice. The role of H-2D, H-2K, and non-H-2 antigens in rejection. Transplantation. 1973;16(4):343–50.

89. Galli SJ, and Hammel I. Unequivocal delayed hypersensitivity in mast cell-deficient and beige mice. Science. 1984;226(4675):710-3.

90. Singer M, Wang C, Cong L, Marjanovic ND, Kowalczyk MS, Zhang H, et al. A Distinct Gene Module for Dysfunction Uncoupled from Activation in Tumor-Infiltrating T Cells. Cell. 2016;166(6):1500–11 e9.

