## Supplemental Material for "Neuropilin-2 functions as a co-inhibitory receptor to regulate antigen-induced inflammation and allograft rejection"

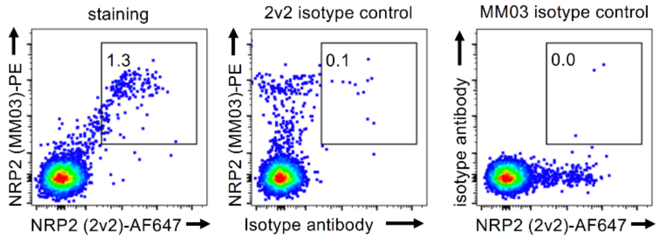


**Supplemental Figure 1:** Specificity of NRP2 flow cytometric staining was evaluated using freshly isolated PBMC and isotype control antibodies. Dot plots show $CD4^+$ T cells (gated).

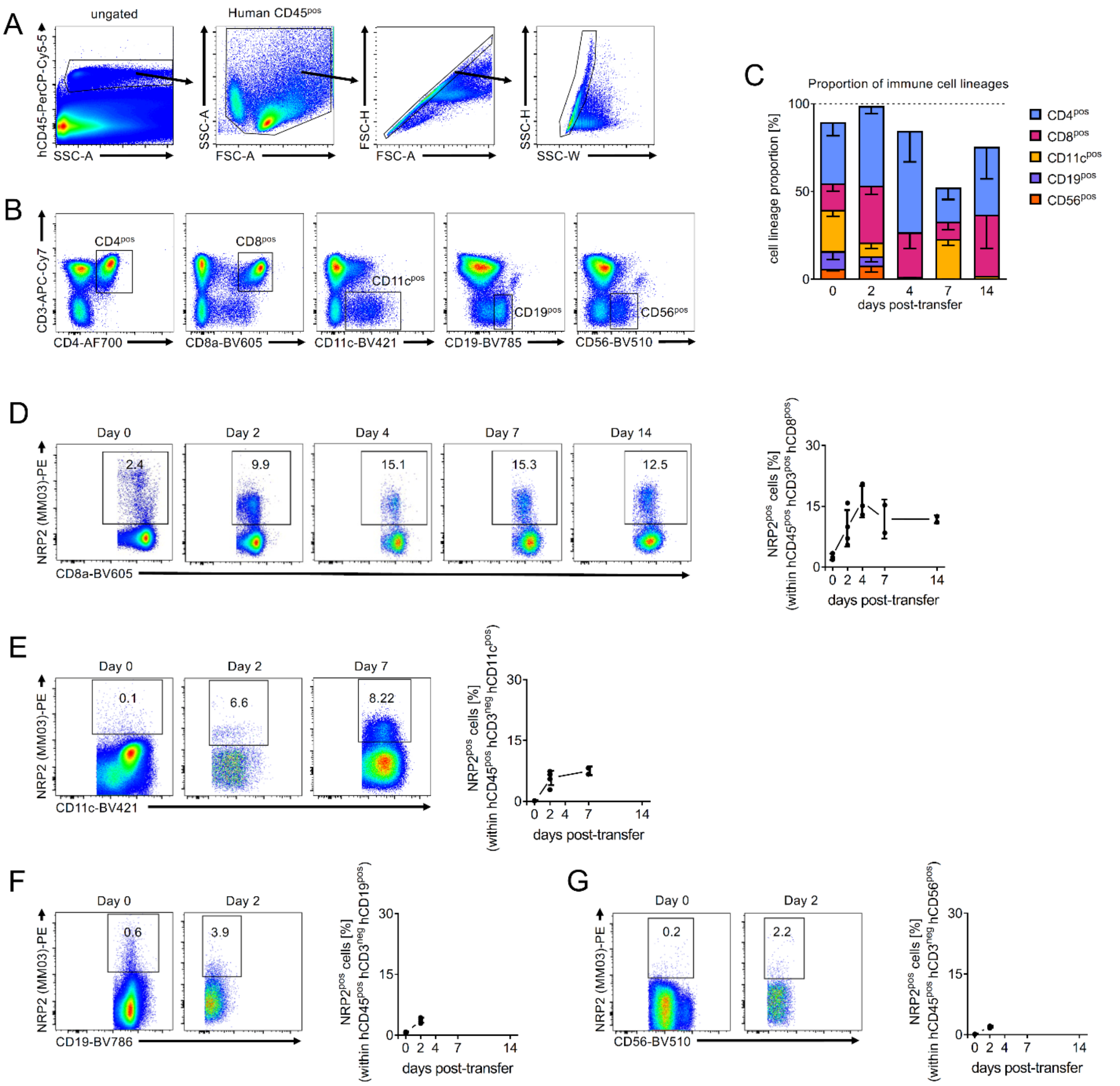


**Supplemental Figure 2: Analysis of NRP2 expression on human immune cells within huSCID mice. (A)** Gating stragtegy for the identification of human cells from huSCID mice (also depicted in Figure 1F-G). **(B)** Identification of human immune cell lineages isolated from splenocytes of huSCID mice. Dotplots are gated on human CD45$^{pos}$ cells. **(C)** Bar graph represents the percent of each lineage and lineage perisitence within human CD45$^{pos}$ cells at different time-points post-transfer in SCID-beige mice. NRP2 expression on **(D)** human CD8$^{+}$ T cells, **(E)** human CD11c$^{+}$ myeloid cells, **(F)** human CD19$^{+}$ B cells, and **(G)** human CD56$^{+}$ NK cells for up to 14 days dependent on survivial within the huSCID. Representative dot plots are gated on human CD45$^{+}$ and each lineage marker$^{+}$. Line graphs illustrate a summary of NRP2 expression in n=2-4 independent experiments per time-point.

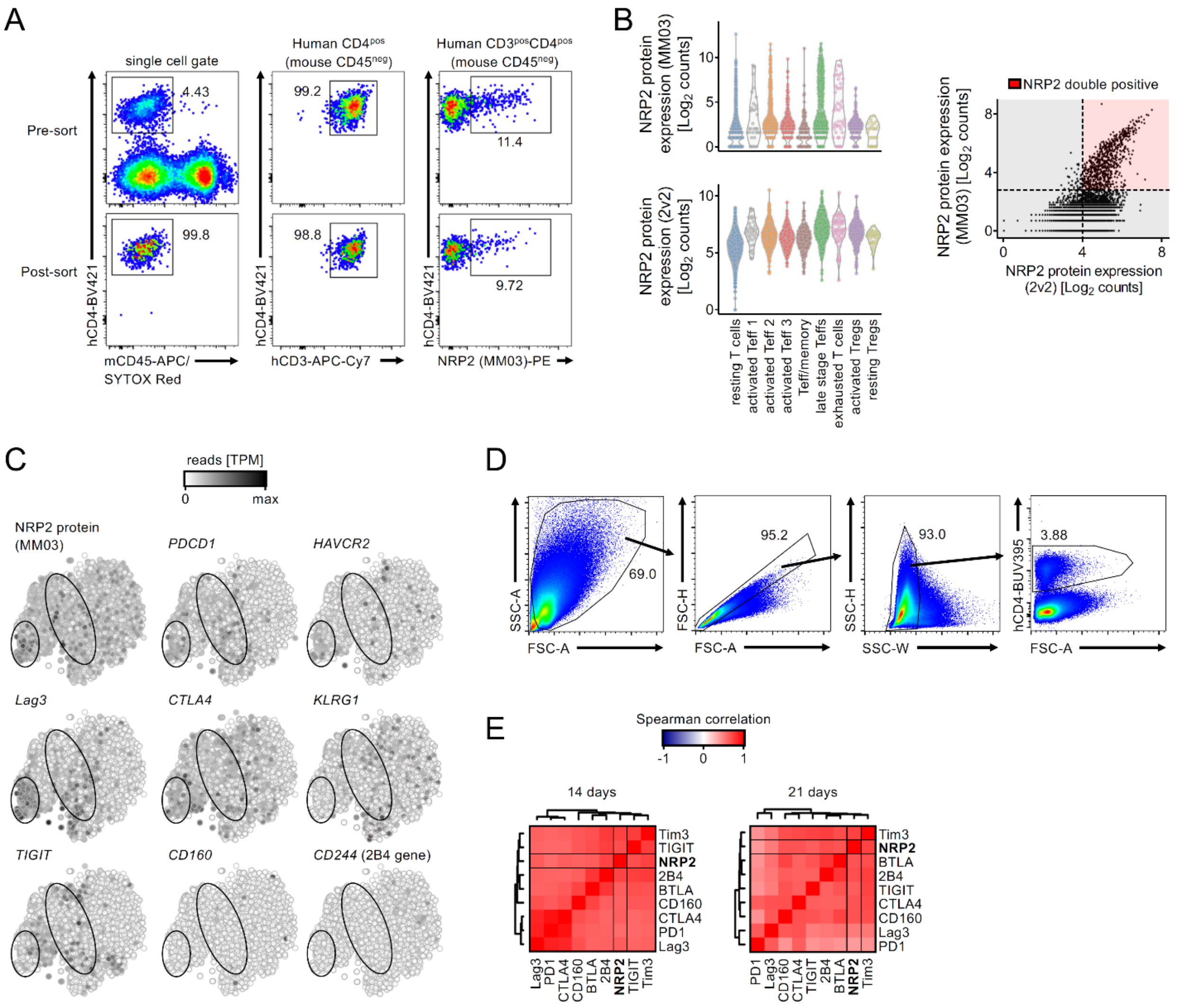


**Supplemental Figure 3: Gating strategies and raw data used in Figure 2. (A)** Sorting strategy and purity control for sorting of human CD3$^{+}$CD4$^{+}$ T cells from the huSCID model on day 7 post transfer (mCD45: mouse CD45; hCD3: human CD3; hCD4: human CD4). Single cell gating, and hCD4 cells in which mCD45 is gated out (CD45$^{neg}$) are shown. **(B)** Left: Violin blots of sequenced NRP2 protein expression on the cells identified in Panel A using CITE-seq (top: clone MM03; bottom: clone 2v2). Right: Scatter plot of scRNA-seq NRP2 antibody counts. Color highlighted areas define NRP2$^{pos}$ and NRP2$^{neg}$ cells for each antibody. Positivity using both antibodies is highlighted in red. **(C)** tSNE plots (grey-scale highlighted) of co-inhibitory receptor mRNA expression and NRP2 protein expression (using the MM03 clone). NRP2 enriched regions are highlighted by circles (see Figure 2A-B). **(D)** Gating strategy for flow cytometric analysis of human CD4$^{+}$ T cells used in Figure 2E-J. **(E)** Flow cytometric analysis was performed on CD4$^{+}$ T cells harvested from the huSCID model on day 7 (Figure 2E), day 14 and day 21 post humanization. Heatmaps were generated to illustrate the mean spearman's rank correlation coefficient between the expression of NRP2 and co-inhibitory receptors (n=4 independent experiments per time-point).

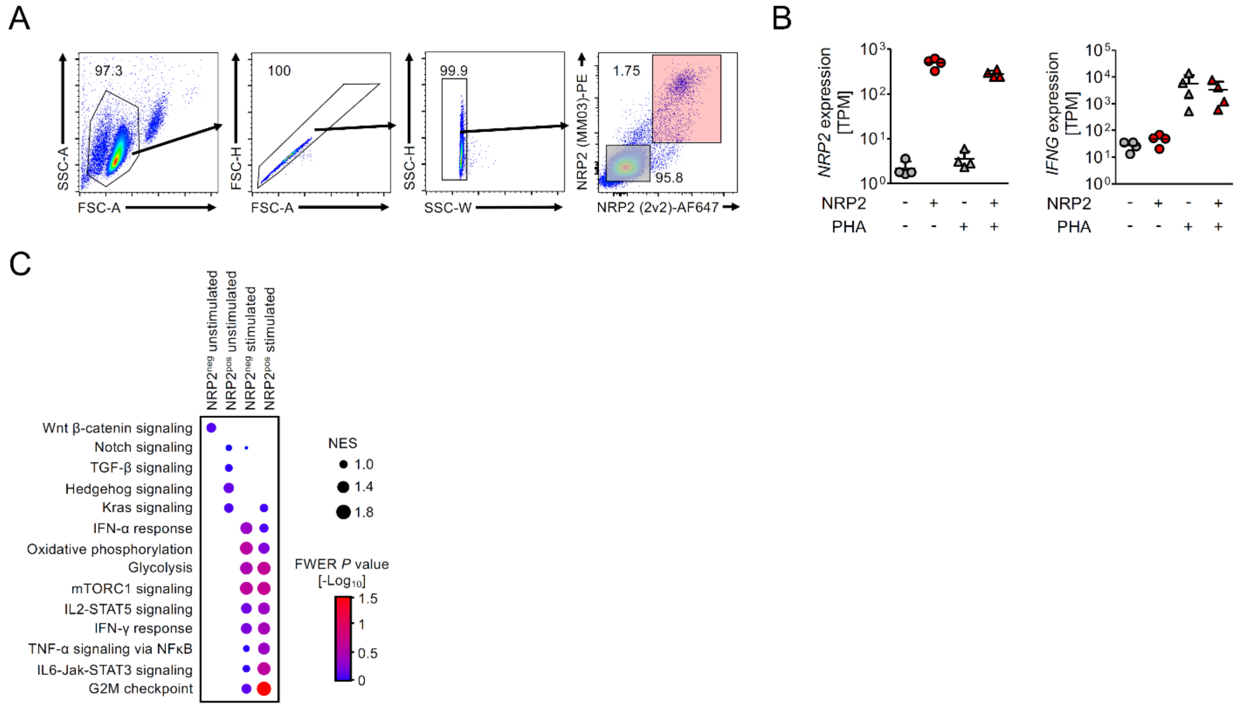


**Supplemental Figure 4: Gating strategy and extended analysis of bulk RNA-seq data. (A)** Sorting strategy for the purification of NRP2$^{pos}$ and NRP2$^{neg}$ cells. **(B)** *NRP2* and *IFNG* mRNA expression in different conditions as quality controls for sorting and mitogen activation respectively. **(C)** Dot plot representing normalized enrichment scores (NES) and familywise error rate adjusted (FWER) *P* values for the gene set enrichment analysis. Signaling pathway activities of cells from four conditions NRP2$^{pos}$ and NRP2$^{neg}$ cells either unstimulated or PHA-stimulated (3 µg/ml for 16 hours) was estimated using the Hallmark gene set database.

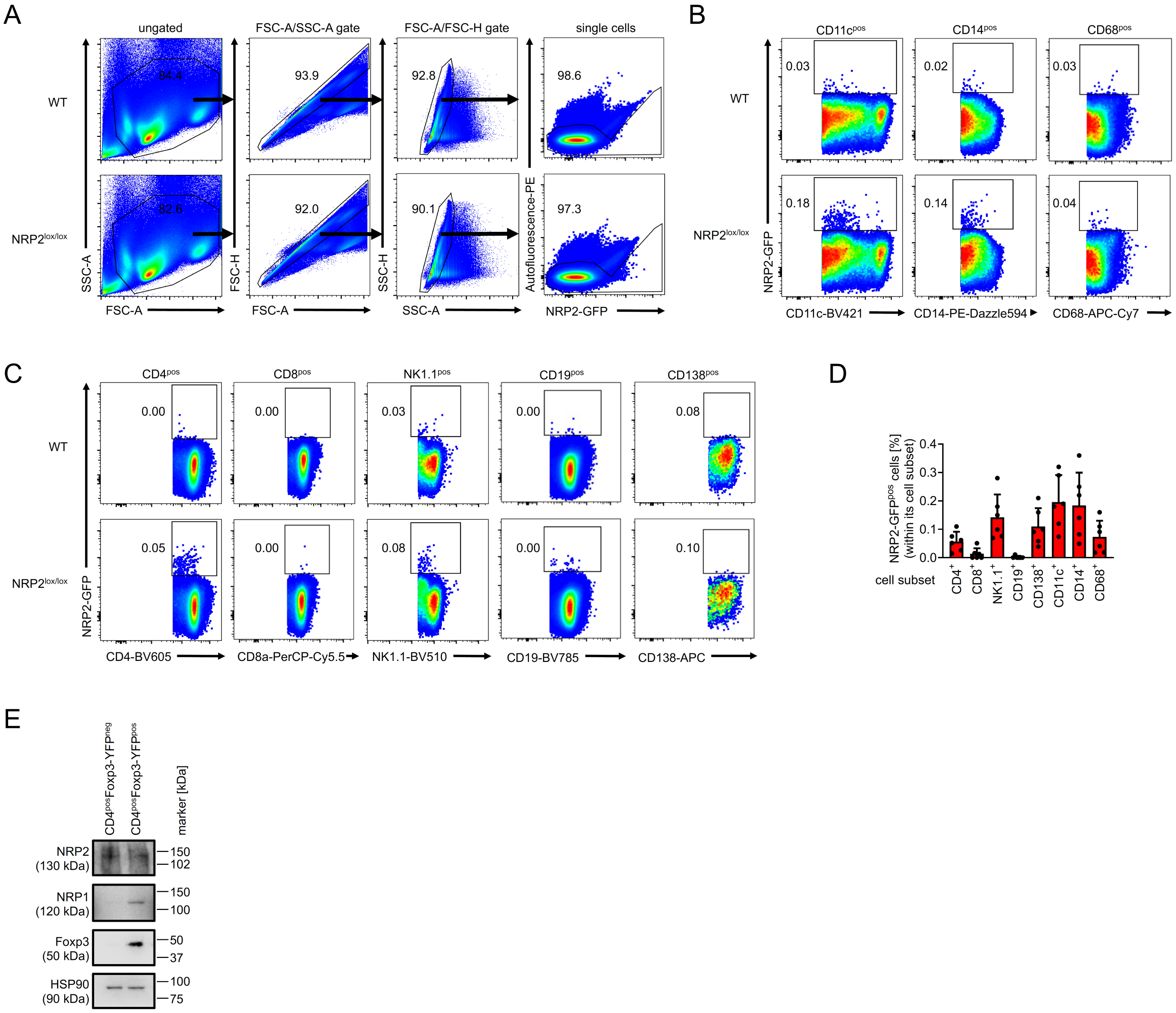


**Supplemental Figure 5: NRP2 expression within murine splenocytes. (A)** Gating strategy to identify NRP2-GFP expression in splenocytes. WT C57BL/6 mice were used to distinguish autofluoresence from specific NRP2-GFP signals in NRP2$^{lox/lox}$ mice. **(B-C)** Representative dot plots and **(D)** a summary of n=6 independent experiments of NRP2-GFP expression within splenocyte subsets. **(E)** Western blot analysis of FACS-sorted CD4$^{+}$Foxp3-YFP$^{pos}$ (Treg) and CD4$^{+}$Foxp3-YFP$^{neg}$ (Teff) cells isolated from splenocytes of Foxp3-cre mice. Representative of n=4 independent experiments.

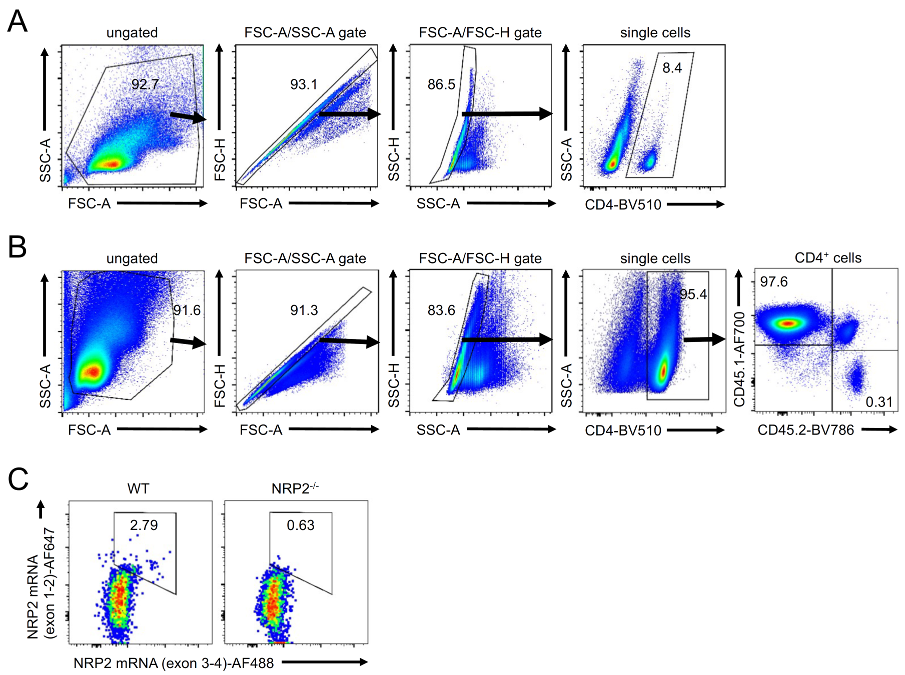


**Supplemental Figure 6: Gating strategy and controls for PrimeFlow assays. (A)** Gating strategy to identify CD4$^{+}$ T cells depicted in Figure 4D-F. **(B)** Gating strategy to identify CD45.1$^{pos}$ CD45.2$^{neg}$ host and CD45.1$^{pos}$ CD45.2$^{pos}$ OT-II CD4$^{+}$ T cells (depicted in Figure 4G-J). **(C)** Splenocytes from C57BL/6 WT and NRP2$^{-/-}$ were stained in the same experiment to identify the gate for NRP2 mRNA positivity. Dot plots show the positive (left: WT splenocytes) and negative (right: NRP2 KO splenocytes) technical controls for the fluorescence in situ hybridization.

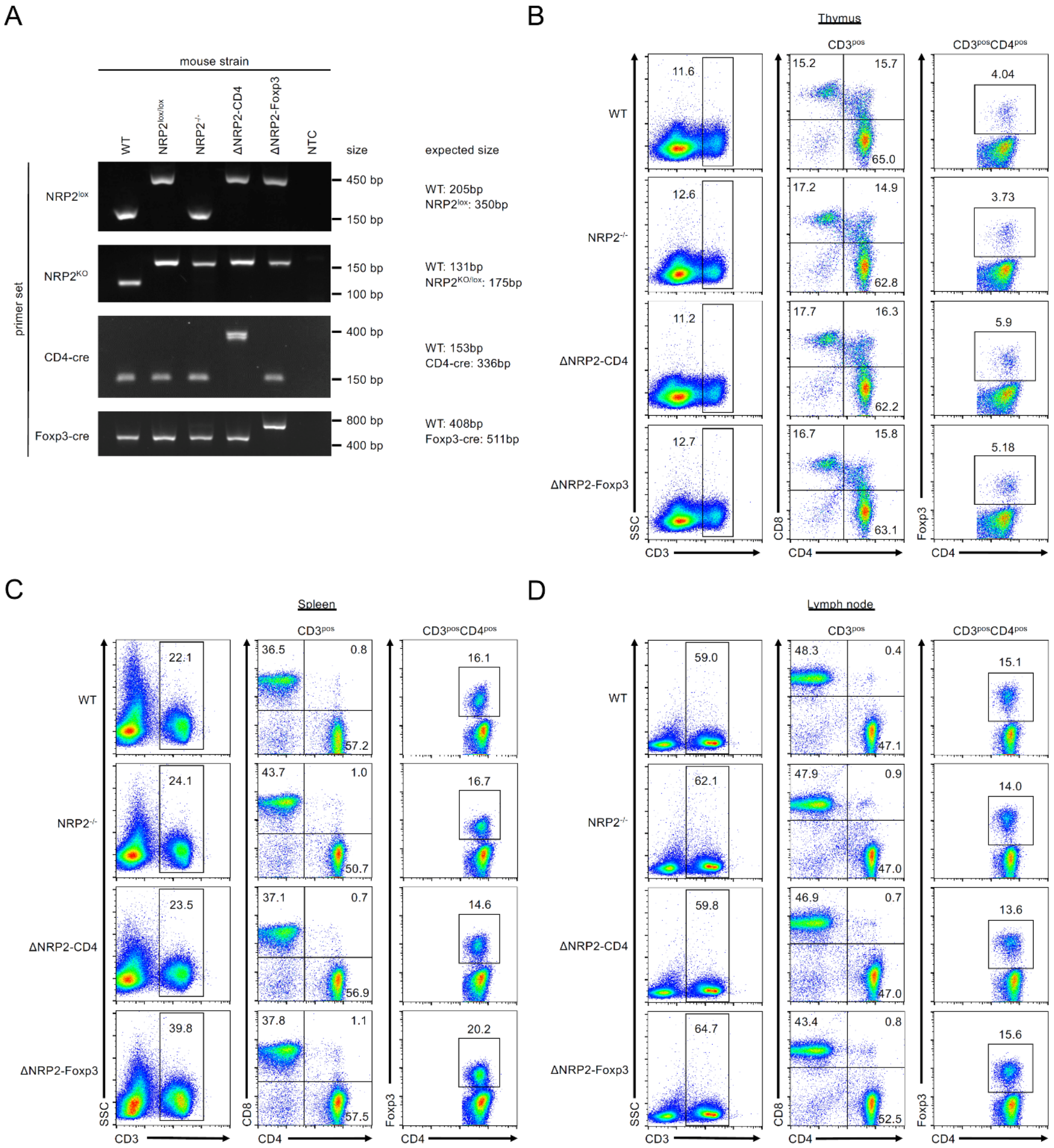


**Supplemental Figure 7: Phenotyping of transgenic mice. (A)** Genotyping of C57BL/6 WT, global NRP2$^{-/-}$ knockout, conditional ΔNRP2-CD4 knockout and ΔNRP2-Foxp3 knockout mice (DNA from tail biopsies). **(B-D)** Immunophenotyping of WT and transgenic mice by flow cytometry. Dot plots are representative of n=3 mice initially evaluated. NRP2$^{lox/lox}$ and Foxp3-cre mice were similar to WT mice (not shown). Frequencies of CD3, CD4, CD8 and Foxp3 positive cells in the **(B)** thymus **(C)** spleen and **(D)** inguinal lymph nodes.

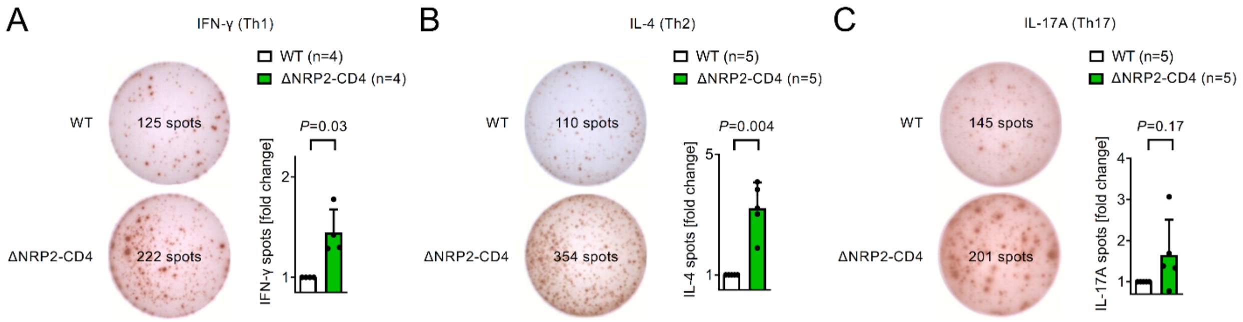


**Supplemental Figure 8: Knockout of NRP2 in murine CD4$^+$ T cells increases T helper cell differentiation *in vitro*.** Naive CD4$^+$ T cells from either WT or ΔNRP2-CD4 knockout mice were cultured in T helper cell (Th) polarizing conditions for 48 hours as described (78), rested for an additional 48 hours, and **(A)** Th1, **(B)** Th2 and **(C)** Th17 differentiation was evaluated using ELISPOT assays respectively. Representative images of ELISPOT wells (left) and bar graphs (from n=4-5 experiments, right) summarizing the fold change in IFN-γ, IL4 and IL-17A comparing knockout cells to WT cells (fold change of mean spots/well ± SD; One sample t-test).

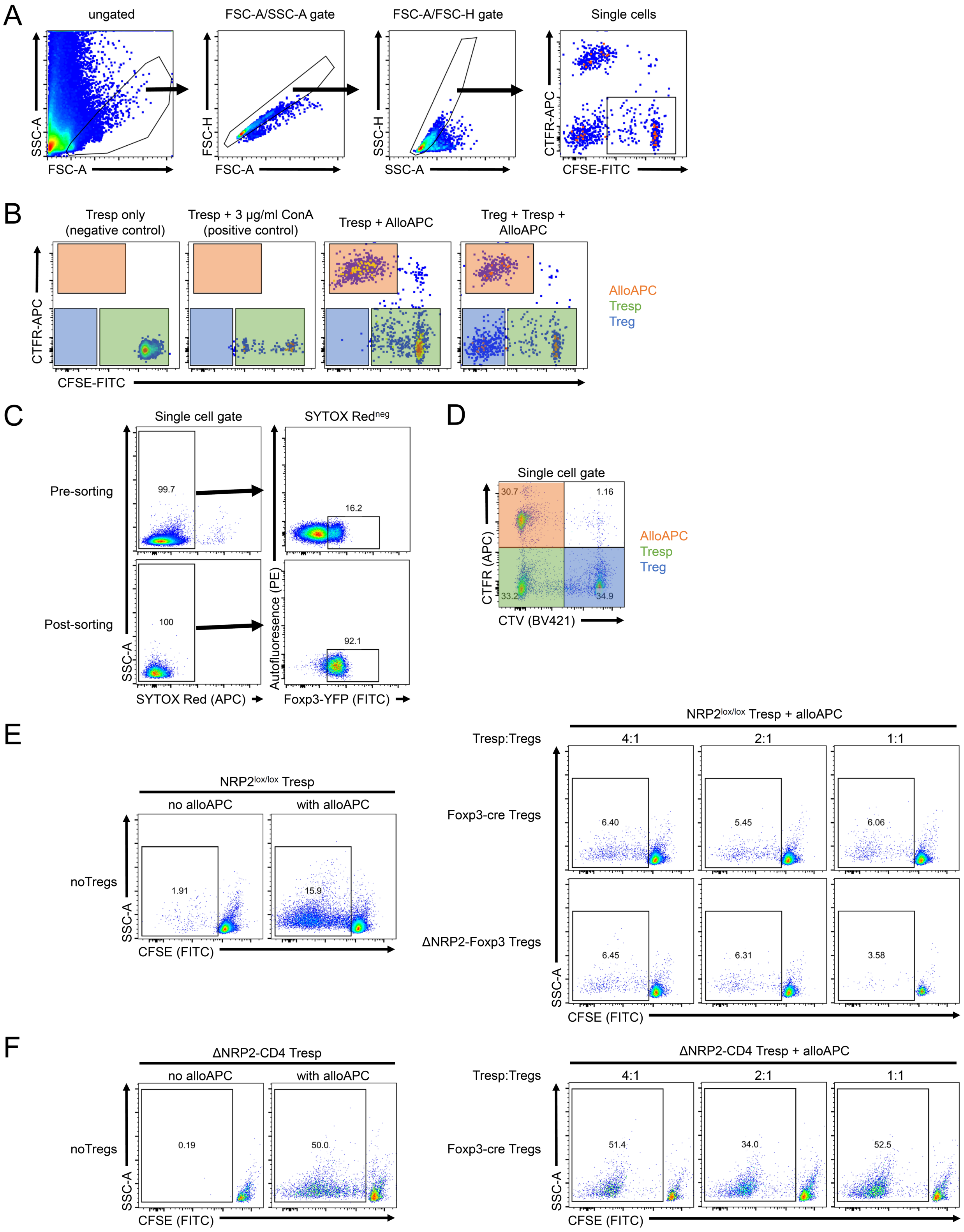


**Supplemental Figure 9: Gating strategy and controls for Treg suppression assays.** C57BL/6 $NRP2^{lox/lox}$ and ΔNRP2-CD4 knockout mice received a fully MHC mismatched Balb/c skin transplant; Teff and Treg were harvested on day 14 and in vitro suppression assays were performed following coculture with irradiated Balb/c APCs as in Figure 6G-I. **(A)** Gating strategy and **(B)** controls to identify a Tresp gate (highlighted in green). The same gating strategy and controls were used for Treg suppression assays in Figure 6G-H. **(C-F)** Validation in vitro suppression experiment using purified populations of Foxp3-YFP Tregs (complimentary to data shown in Figure 6G-I). C57BL/6 NRP2$^{lox/lox}$, ΔNRP2-CD4, Foxp3-cre and ΔNRP2-Foxp3 mice received a fully MHC mismatched Balb/c skin transplant. On day 14 post-transplant, CD4$^{+}$Foxp3-YFP$^{+}$ Tregs were FACS-sorted from Foxp3-cre and ΔNRP2-Foxp3 mice and stained with CellTrace Violet (CTV). CD4$^{+}$CD25$^{neg}$ Tresp were harvested from NRP2$^{lox/lox}$ and ΔNRP2-CD4 recipients and stained with CFSE. Treg function was assessed in an in vitro suppression assay as depicted in Figure 6G-I using irradiated and CellTrace Far Red (CTFR)-stained Balb/c splenocytes (alloAPC; Tresp:alloAPC ratio 1:1). **(C)** Dotplots depict cell viability and Foxp3 positivity pre- and post-sorting. **(D)** Representative dotplot illustrates the gating strategy to identify CTV$^{neg}$CFSE$^{pos}$CTFR$^{neg}$ Tresp cells for evaluating Tresp proliferation using CFSE dilution. **(E-F)** Proliferation of NRP2 WT (E) and NRP2 KO (F) Tresp without Tregs (left panels) or with increasing ratios of either NRP2 WT (right panel; top row) of KO (right panel; bottom row) Tregs. One validation experiment was performed.

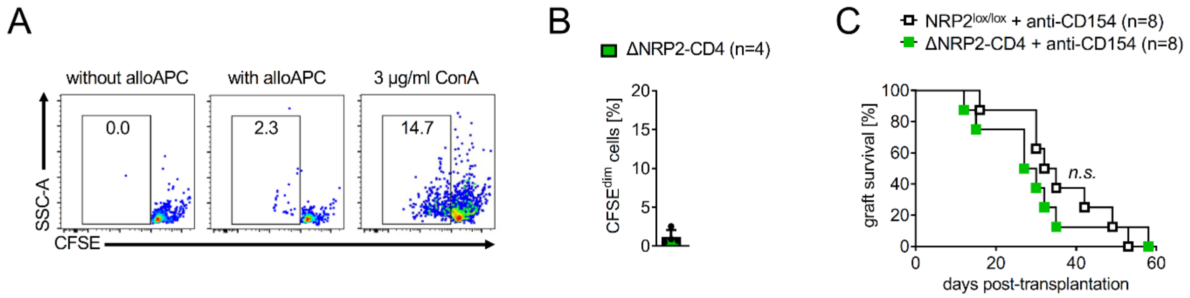


**Supplemental Figure 10: Costimulatory blockade using anti-CD154 inhibits allogeneic priming of Teffs and abrogates NRP2 function. (A-B)** Fully MHC mismatched Balb/c hearts were transplanted into NRP2$^{lox/lox}$ wild type or ΔNRP2-CD4 knockout recipients treated with anti-CD154 on days 0 and 2 post transplantation. Allogeneic priming on day 14 post-transplantation was evaluated by coculture of recipient CD4$^{+}$ T cells with irradiated donor Balb/c splenocytes for 5 days; proliferation as assessed by CFSE dilution. Concavalin A (ConA, 3 µg/ml) was used as a positive control. **(A)** Representative dot plots and **(B)** bar graph of n=4 mice is depicted (frequency of alloreactive ΔNRP2-CD4 knockout CFSE$^{dim}$ cells ± SD)**. (C)** Kaplan-Meier graft survival curves (*n.s.* not significant).

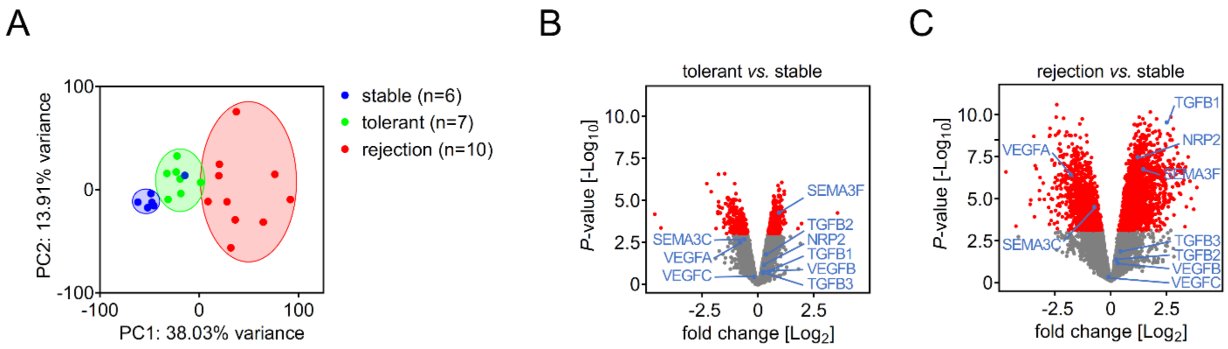


**Supplemental Figure 11: Expression of NRP2 ligands in tolerant renal transplant biopsies.** We analyzed a published database of renal transplant biopsies (GSE106675) that were taken from renal transplant recipients following tolerance induction using the Northwestern protocol (76-77). The database also included findings from surveillance biopsies from a control cohort of stable transplants recipients on conventional immunosuppression, and from patients with clinical rejection. **(A)** Principal component analysis of all samples. **(B-C)** Volcano plot analysis comparing stable patients on immunosuppression to **(B)** biopsies from tolerant patients off immunosuppression or **(C)** biospies with evidence of rejection. Statistically significant ($P$<0.001) regulated transcripts are highlighted in red, and NRP2 ligands are highlighted as blue dots.

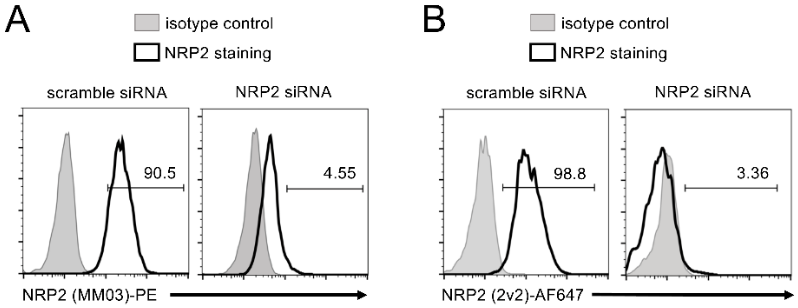


**Supplemental Figure 12: NRP2 antibody specificity.** The specificity of the two anti-human NRP2 antibody clones used in this study was assessed in a knockdown approach using the NRP2-expressing U87MG glioblastoma cell line as previously described (45). **(A)** Binding of the MM03 anti-NRP2 clone, and **(B)** binding of the 2v2 anti-NRP2 clone to scramble siRNA (left panels) or NRP2 siRNA (right panels) transfected U87MG cells.

**Supplemental Table 1. Antibodies used in these studies.**

| target (fluorochrome/enzyme/DNA-oligo) | clone | company |
|---|---|---|
| anti-Biotin (A0436 [DNA]) | 1D4-C5 | Biolegend |
| anti-BrdU (AF488, AF647) | 3D4 | Biolegend |
| anti-human 2B4 (AF700) | C1.7 | Biolegend |
| anti-human BTLA (PE-Cy5) | MIH26 | Biolegend |
| anti-human CD11c (BV421) | 3-HCL-3 | Biolegend |
| anti-human CD160 (PE) | 51-10C9 | BD Biosciences |
| anti-human CD19 (BV785) | HIB19 | Biolegend |
| anti-human CD279/PD-1 (BV421) | NAT105 | Biolegend |
| anti-human CD3 (unconjugated, APC-Cy7) | HIT3a | Biolegend |
| anti-human CD38 (BV605) | HB-7 | Biolegend |
| anti-human CD366/Tim3 (APC-Cy7) | F38-2E2 | Biolegend |
| anti-human CD4 (BUV395) | RPA-T4 | BD Biosciences |
| anti-human CD4 (BV421, APC, AF700) | OKT4 | Biolegend |
| anti-human CD45 (PerCP-Cy5.5) | HI30 | Biolegend |
| anti-human CD56 (BV510) | 5.1H11 | Biolegend |
| anti-human CD69 (BV421) | FN50 | Biolegend |
| anti-human CD8a (BV605) | HIT8a | Biolegend |
| anti-human CTLA4 (BV605) | BNI3 | Biolegend |
| anti-human HLA-DR (AF488) | L243 | Biolegend |
| anti-human IFNγ (AF488) | 4S.B3 | Biolegend |
| anti-human Lag3 (BV510) | 11C3C65 | Biolegend |
| anti-human NRP2 (biotin, AF647) | 2v2 | aTyr Pharma |
| anti-human NRP2 (unconjugated, PE, APC) | MM03 | Sino Biologics |
| anti-human TIGIT (PE-Cy7) | A15153G | Biolegend |
| anti-human TruStain FcX | --- | Biolegend |
| anti-mouse CD11c (BV421) | N418 | Biolegend |
| anti-mouse CD138/Syndecan-1 (APC) | 281-2 | Biolegend |
| anti-mouse CD14 (PE-Dazzle594) | Sa14-2 | Biolegend |
| anti-mouse CD19 (BV785) | 6D5 | Biolegend |
| anti-mouse CD197/CCR7 (BV605) | 4B12 | Biolegend |
| anti-mouse CD279/PD1 (BV785, PerCP-Cy5.5) | 29F.1A12 | Biolegend |
| anti-mouse CD3 (PE) | 17A2 | Biolegend |
| anti-mouse CD366/Tim3 (PE, APC-Fire750) | B8.2C12 | Biolegend |
| anti-mouse CD4 (BV510, BV605, BV785, PerCP-Cy5.5, AF700) | GK1.5 | Biolegend |
| anti-mouse CD44 (AF488, PerCP-Cy5.5) | IM7 | Biolegend |
| anti-mouse CD45 (APC) | 30-F11 | Biolegend |

| | | |
|---|---|---|
| anti-mouse CD45.1 (BV510, AF700) | A20 | Biolegend |
| anti-mouse CD45.2 (BV786, PE-Dazzle594) | 104 | Biolegend |
| anti-mouse CD62L (BV421, PE) | MEL-14 | Biolegend |
| anti-mouse CD68 (APC-Cy7) | FA-11 | Biolegend |
| anti-mouse CD69 (PE, PE-Dazzle594) | H1.2F3 | Biolegend |
| anti-mouse CD8a (PerCP-Cy5.5, AF647) | 53-6.7 | Biolegend |
| anti-mouse CXCR5 (BV605) | L138D7 | Biolegend |
| anti-mouse Foxp3 (unconjugated, AF647) | FJK-16s | Thermo Fisher |
| anti-mouse HSP90 | C45G5 | Cell Signaling |
| anti-mouse IFNγ | AN-18 | Biolegend |
| anti-mouse IFNγ (biotin) | R4-6A2 | Biolegend |
| anti-mouse IgG (H+L) (HRP) | polyclonal | Vector Laboratories |
| anti-mouse IgG-Fcγ fragement (AF647) | polyclonal | Jackson Immuno |
| anti-mouse IL17A | eBio17CK15A5 | Thermo Fisher |
| anti-mouse IL17A (biotin) | eBio17B7 | Thermo Fisher |
| anti-mouse IL2 | JES6-1A12 | Biolegend |
| anti-mouse IL2 (biotin) | JES6-5H4 | Biolegend |
| anti-mouse IL4 | 11B11 | Biolegend |
| anti-mouse IL4 (biotin) | BVD6-24G2 | Biolegend |
| anti-mouse NK-1.1 (BV510) | PK136 | Biolegend |
| anti-mouse NRP1 | D62C6 | Cell Signaling |
| anti-mouse NRP2 | D39A5 | Cell Signaling |
| anti-mouse TruStain FcX | --- | Biolegend |
| anti-phycoerythrin (A0911 [DNA]) | PE001 | Biolegend |
| Streptavidin (FITC) | polyclonal | Thermo Fisher |

**Supplemental Table 2. Primers used in these studies.**

| target | reverse | forward |
|---|---|---|
| NRP2-WT | 5` CAG GTG ACT GGG GAT AGG GTA 3` | 5` AGC TTT TGC CTC AGG ACC CA 3` |
| NRP2lox | 5` CCT GAC TAC TCC CAG TCA TAG 3` | |
| NRP2-WT | 5` TGA AAT CAG CAA AGA GGG AGC 3` | 5` GGG AAA CCC TCG TGA TGT TGT 3` |
| NRP2$^{KO}$ | 5` ATC TTC CAT CAC GTC GAA CTC 3` | |
| CD4-WT | 5` TAT GCT CTA AGG ACA AGA ATT GAC A 3` | 5` GTT CTT TGT ATA TAT TGA ATG TTA GCC 3` |
| CD4-cre | 5` CTT TGC AGA GGG CTA ACA GC 3` | |
| Foxp3-WT | 5` GTG TGA CTG CAT GAC TAA CTT TGA 3` | 5` CCC TTG ACC TCA AAA CCA AG 3` |
| Foxp3-cre | | 5` TGG CTG GAC CAA TGT GAA C 3` |
